# Community performance curves predict temporal stability despite interaction effects

**DOI:** 10.64898/2026.03.27.714753

**Authors:** Francesco Polazzo, Til Hämmig, Paula Eileen Stöckli, Shyamolina Ghosh, Stephan B. Munch, Owen L. Petchey

## Abstract

Predicting whether aggregate community properties remain stable as environments fluctuate is a central challenge of ecological research. A critical open question is, however, how well the fundamental niche of species can predict this aspect of community stability. We conceptualised community performance (CP) curves by aggregating species-specific performance across environmental conditions and asked whether the variability of these curves explains community temporal variability. In Lotka–Volterra and in mechanistic consumer–resource simulations, CP variability explained on average 92% of the variation in total abundance variability. Explanatory power remained above 75% across gradients of interaction strength and coexistence conditions. Derivation of the expected community variability – CP variability relationship for the Lotka-Volterra case explains this high explanatory power despite interaction effects. In a purpose-designed microcosm experiment, CP variability explained 67% of variation in the temporal variability of total community biomass. Explanatory power of CP variability therefore retains substantial information about community stability even in strongly interacting systems, providing a tractable basis for forecasting stability when species interactions are unknown.

**One-sentence summary:** Aggregated species performance curves predict community stability even when interactions strongly reorganise population synchrony.

**Open Science Statement:** Data and code to reproduce the analysis is available at https://github.com/FrancescoPola/CPCs_predict_CommunityStability, and will be permanently deposited in Zenodo upon publication.

## Introduction

Forecasting whether ecosystems will remain stable as environments fluctuate is a central challenge in ecology. Research on temporal stability typically focuses on aggregate community or ecosystem properties, and quantifies the extent to which total abundance, biomass, or productivity varies through time (Isbell *et al*. 2015; Loreau & de Mazancourt 2013; Pennekamp *et al*. 2018; Thibaut & Connolly 2013; Yachi & Loreau 1999). This framework differs from classical equilibrium notions of stability, which historically focused on whether a system returns to a baseline state after a single perturbation, or whether interacting species can locally coexist (Allesina & Tang 2012; Chesson 2000; May 1973; Pimm 1984). Instead, our focus centers on the stability of an aggregate property in a continuously variable environment, and what properties of a community can result in relatively constant aggregate properties as environmental conditions fluctuate.

Temporal stability is an aggregate property, but it is the result of population level dynamics. Environmental variation first acts through species-specific response functions (Ives *et al*. 1999). When species differ in the conditions that maximise performance, or in the breadth and shape of their responses, the same environmental sequence can drive dissimilar fluctuations among populations. Theory therefore establishes a qualitative link between such response diversity and aggregate biomass variability: heterogeneous responses can reduce synchrony and buffer variation in aggregate community properties even when individual populations are variable (Doak *et al*. 1998; Loreau & de Mazancourt 2008; Yachi & Loreau 1999).

Yet species do not experience environmental fluctuations in isolation. Interspecific interactions can reshape environmental responses (Ntetsika *et al*. 2026), potentially amplifying or dampening environmentally driven fluctuations and altering the underlying covariance structure among populations (Gonzalez & Loreau 2009; Ives *et al*. 1999; McCann *et al*. 1998). This could make predicting stability from species responses challenging. However, the issue is not whether interactions matter; they clearly do. The question is whether the environmental response structure of a community can nonetheless provide a practical and general predictor of temporal stability when detailed information on interaction strengths is unavailable.

Answering this question has been difficult for both historical and methodological reasons. Historically, ecological stability theory has traditionally focused on coexistence, persistence, and return to equilibrium, from classical complexity analyses to modern interaction-network approaches, thereby making interaction structure central (Allesina & Tang 2012; Grilli *et al*. 2016; May 1973). However, aggregate temporal stability questions ask not whether every population persists or returns to equilibrium, or whether there is coexistence, or how robust a network is, but how species-level fluctuations combine into the variability of a total community or ecosystem property. In this context, interaction effects on aggregate temporal stability could be positive or negative. Interspecific competition can generate compensatory dynamics by reducing synchrony among species, but can also increase population-level variability (Ives *et al*. 1999; Loreau & de Mazancourt 2013). Depending on interaction symmetry (i.e. how similarly species affect one another) and strength, these effects can cancel or produce either stabilizing or destabilizing net effects on community variability (Hughes & Roughgarden 1998; Loreau & de Mazancourt 2008).

Empirically, however, this possibility has rarely been tested by placing environmental responses and interactions within the same predictive framework. Our literature review found that both mechanisms are widely invoked, but seldom compared within the same framework (Supplementary Information 1). Moreover, while theory and experiments increasingly find that differential environmental responses (i.e., response diversity) and species-specific dynamics can stabilize aggregate properties (Danet *et al*. 2025; de Mazancourt *et al*. 2013; Meng *et al*. 2025; Polazzo *et al*. 2025; Ruiz-Moreno *et al*. 2024), experts continue to rank interactions among the principal obstacles to using response diversity predictively (Ross *et al*. 2026).

The second obstacle is methodological. Environmental responses are commonly represented by single traits or moments of trait distributions, such as a community-weighted mean or variance. These summaries identify where species perform best but discard the intrinsic non-linearity of species performance-environment relationships. What is missing is response shape. A thermal optimum, for example, tells us where a species performs best. It does not tell us, however, how rapidly performance declines away from that optimum, whether tolerance is broad or narrow, whether the curve is symmetric or skewed, or whether different species’ curves complement one another (Angilletta 2006; Arnoldi *et al*. 2025). Communities with identical means and variances in thermal optima can nonetheless differ sharply in response breadth, skew, overlap, and complementarity. Because performance–environment relationships are nonlinear, those differences determine how environmental variation is converted into temporal variation (Angilletta 2006; Dell *et al*. 2011; Huey & Kingsolver 1989).

We address this limitation by treating each species performance curve as a metatrait: integrated response profiles that capture how multiple underlying traits jointly determine performance across an environmental gradient. We then aggregate these species-level curves into a *community performance (CP) curve*, which describes the expected performance of a community across environmental conditions. Evaluating a CP curve over an environmental sequence produces a distribution of expected community performance (Fig. 1). Its coefficient of variation is an explicit prediction of temporal variability of total community performance: flat CP curves predict relatively constant aggregate performance, whereas strongly varying CP curves predict larger fluctuations (just as would be the case for species performance curves and fluctuation in population abundance).

**Figure 1.**
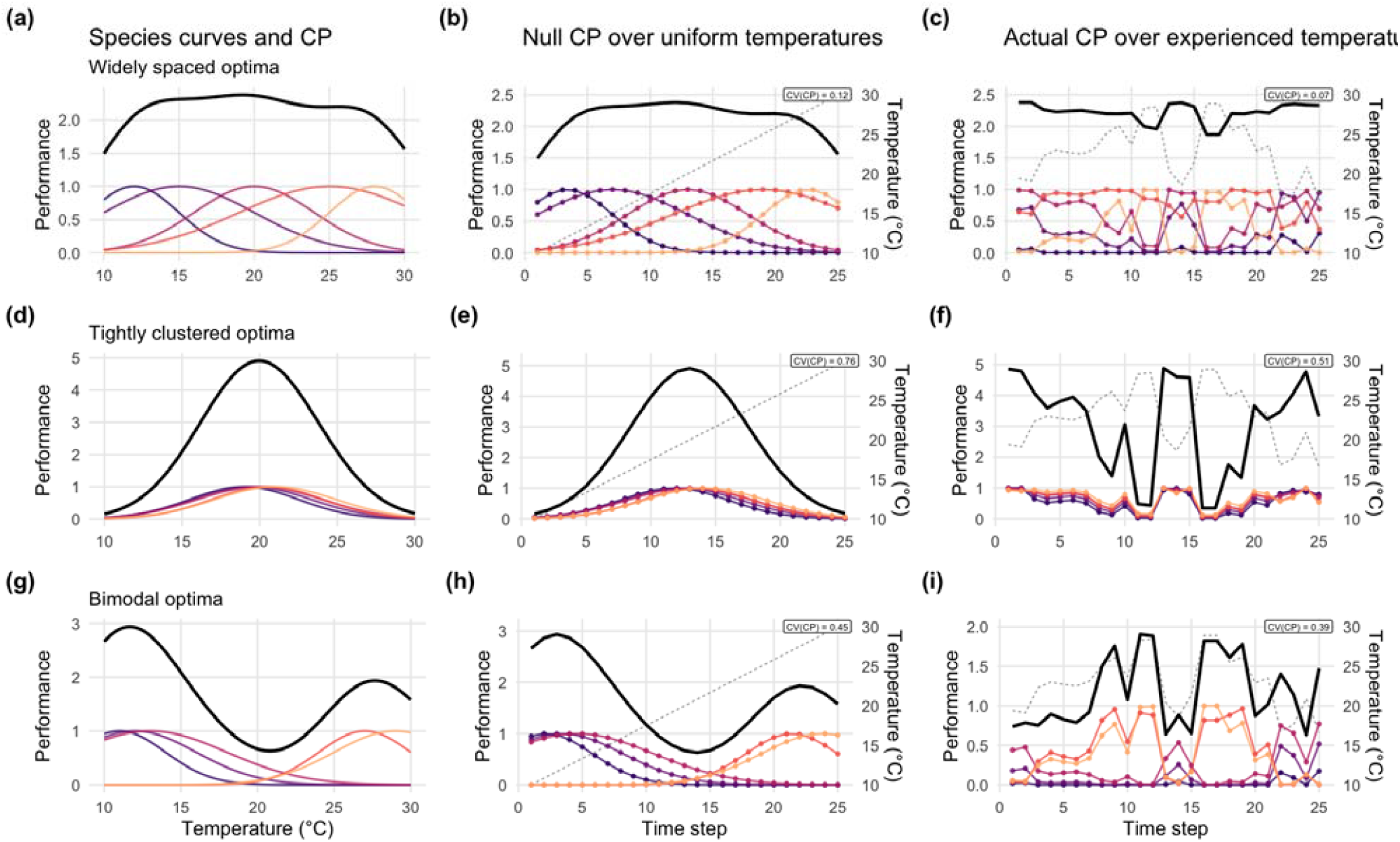
Community performance curves, species performance curves, and temperature driven community performance distributions across four community types. Three idealised community configurations are shown, with species thermal optima that are (a–c) widely spaced, (d–f) tightly clustered, (g–i) skewed toward cooler temperatures. In the first column, thin coloured lines show species-level thermal performance curves and thick black lines show their sum, the community performance (CP) curve. In the second column, the same species curves are evaluated over a null temperat re sequence that samples the full thermal range uniformly, producing a null CP time series and its coefficient of variation, CV(CP). In the third column, the curves are evaluated over a realised temperature sequence, producing the actual CP time series and CV(CP). In the time-series panels, coloured lines and points show species-level performance, thick black lines show summed community performance, and dashed grey lines show the corresponding temperature sequence. These examples illustrate that CP variability is determined jointly by the arrangement of species response curves and by the environmental conditions the community experiences.

CP variability can be calculated using the realised environmental sequence a community experiences (*actual* CP variability) or, when that sequence is unavailable, a null assumption, such as a uniform distribution across the relevant environmental range (*null* CP variability). The framework therefore separates information contained in the CP curve shape from information added by the environmental frequency distribution (Fig. 1). It also yields a test of the importance of interaction effects. If the presence of interactions is important, CP-based prediction should deteriorate as interaction effects increase.

Here, we test if CP variability explains variation in community stability using mathematical simulations of communities and a purpose-designed microcosm experiment. A continuous-time Lotka–Volterra model and a mechanistic consumer–resource model independently vary response structure and interaction strength. In parallel, we use thermal performance curves measured in ciliate monocultures to design multispecies communities spanning a predicted gradient of CP variability, then expose them to fluctuating temperature. Together, these approaches ask whether aggregate temporal stability can be predicted from species environmental responses even when detailed interaction matrices are unknown.

## Methods

### Lotka–Volterra simulations

In the Lotka–Volterra simulations, species biomass was modelled in continuous time. The rate of change in the biomass of species depended on its temperature-dependent intrinsic growth rate, its temperature-dependent carrying capacity, the competitive effects of all species in the community, and a constant immigration flux:

Here, *N*_*i*_(t) is the biomass of species *i* at time *t, S* is species richness, α_*ij*_ is the effect of species *j* on species *i, K*_*i*_(*T*_*t*_) is the temperature-dependent carrying capacity of species *i* in the absence of interspecific effects, and 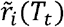 is its temperature-dependent intrinsic growth rate. The implementation defined

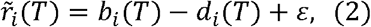

where *ε* = 10^−6^ is a small numerical offset, and set carrying capacity to

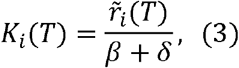

with *β* = *δ* = 0.001. Immigration was continuous: *I* = 0.1 biomass units were added per species per unit time. Thus, 0.1 enters the right-hand side of the differential equation as a constant flux rather than as a discrete pulse after each time step.

Temperature dependence was represented by a Gaussian birth-rate function and an exponential death-rate function:

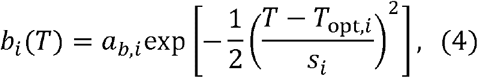

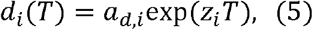

where *a*_*b,i*_ is the birth rate at the species’ thermal optimum *T*_*opt,i*_, *s*_*i*_ controls the breadth of the birth-rate curve, *a*_*d,i*_ is the death rate at *T* = 0, and *z*_*i*_ controls the temperature dependence of death. In the simulations, *a*_*b,i*_ = 0.3 and *a*_*d,i*_ = 0 for all species, making *z*_*i*_ irrelevant.

Communities were assembled using the same factorial design in two independent Lotka–Volterra runs. Species richness was 4, 8, or 12. Species thermal optima were drawn from uniform distributions with means of 16, 20, or 24 °C and ranges of 0, 2, or 4 °C. Thermal breadths were drawn from uniform distributions with mean breadths of 6, 10, or 14 and breadth ranges of 4 or 10. Intraspecific competition was fixed at *α*_*ii*_ = 1. In one run, interspecific competition coefficients were drawn from asymmetric uniform distributions with mean strengths of 0, 0.25, 0.5, 0.75, or 1. In the other, coefficients were drawn from symmetric log-normal distributions with the same median strengths and a log-scale standard deviation of 0.5. Three community replicates were crossed with three temperature time series for each factorial combination, yielding 7,290 simulations per run.

The ordinary differential equations were integrated with the lsoda solver in the deSolve R package (Soetaert *et al*. 2010). Temperature was linearly interpolated during integration. We used relative and absolute error tolerances of 10^−2^ and 10^−4^, respectively, and constrained the maximum integration step to one time unit. Each simulation used a 200-time-unit burn-in followed by 2,000 time units of environmental forcing. During the forced period, temperature had a mean of 20 °C, a standard deviation of 4 °C, and 1/*f* temporal autocorrelation with exponent *γ* = 0.8.

### Consumer–resource simulations

A consumer–resource model (Gonzalez & Feo 2007; Lehman & Tilman 2000) was used as a mechanistic test of CP construction when species abundance is mediated by explicit resource dynamics rather than by a direct Lotka–Volterra performance-to-abundance link. For a community with consumer species and resources, consumer abundance is denoted *N*_*i*_ (*t*), and resource concentration is denoted *R*_*j*_(*t*)□. Consumer dynamics followed

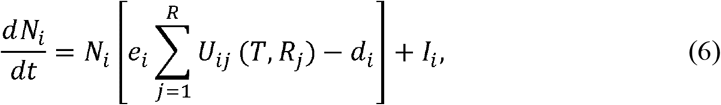

where *e*_*i*_ is conversion efficiency, *d*_*i*_ is mortality, *I*_*i*_ is immigration, and *U*_*ij*_(*T, R*_*j*_) is the uptake rate of consumer i on resource j. Resource dynamics followed

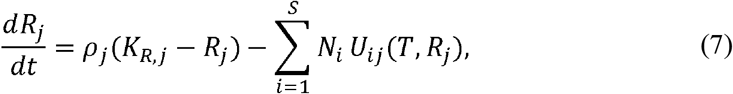

where *ρ*_*j*_ is the resource renewal rate and *K*_*R,j*_ is the resource supply concentration. This chemostat-like renewal term keeps resources bounded in the absence of consumers.

Resource uptake was modelled with Monod kinetics:

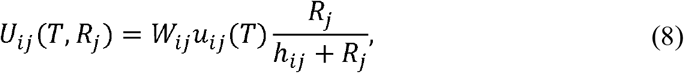

where *W*_*ij*_ is the resource-use weight, *h*_*ij*_ is the half-saturation constant, and *u*_*ij*_(*T*) is the temperature-dependent maximum uptake rate. Temperature dependence of maximum uptake followed a Gaussian curve:

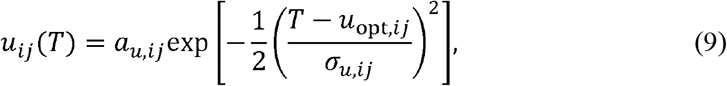

where *a*_*u,ij*_ is the maximum uptake rate, *u*_*opt,ij*_ is the thermal optimum of uptake, and *σ*_*u,ij*_ controls thermal breadth.

The model used a shared-to-private resource-use structure. One resource was shared by all consumers, and each consumer also had one private resource. A specialisation parameter *λ* controlled the resource-use weights:

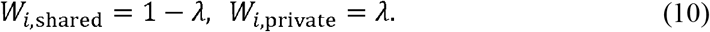

Thus, *λ* = 0 gives complete overlap in resource use, whereas *λ* = 1 gives one private resource per consumer. Intermediate values split uptake between the shared and private resources while holding total resource-use weight constant.

For the consumer–resource model, per-capita growth at temperature *T*, evaluated at resource supply concentrations, was

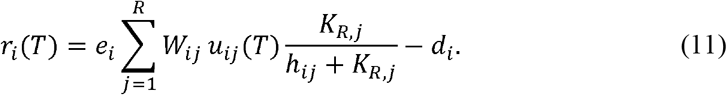

The consumer–resource factorial design paralleled the Lotka–Volterra design. Consumer richness was 4, 8, or 12; mean thermal optimum was 16, 20, or 24 °C; the range of optima was 0, 2, or 4 °C; and mean thermal breadth was 3, 6, and 9, and uptake-breadth range was 0.5 or 1.25. Resource use varied along a private-to-shared gradient. In the deterministic resource-use run, resource-sharing levels were 0, 0.25, 0.5, 0.75, and 1. In the stochastic resource-use run, resource-use weights were drawn from beta distributions centred on the same gradient, with endpoint means adjusted to 0.01 and 0.99 to avoid degenerate beta distributions. Three community replicates and three temperature time series were crossed with each factorial combination, yielding 7,290 valid simulations per run.

Consumer–resource dynamics were integrated with the lsoda solver in the deSolve R package (Soetaert *et al*. 2010). Each simulation used a 200-time-unit burn-in followed by 2,000 time units of environmental forcing. During the forced period, temperature had a mean of 20 °C, a standard deviation of 4 °C, and 1/*f* temporal autocorrelation with exponent *γ* = 0.8. Because uptake rates are temperature dependent and consumers compete through shared resources, the consumer–resource model provides a mechanistic case in which effective interaction strengths can vary with temperature (Duan et al. 2025).

### Community performance

For the Lotka–Volterra model and the empirical microcosm experiment, species performance at temperature *T* was defined as intrinsic growth rate. Community performance was calculated as the sum of species’ performance curves:

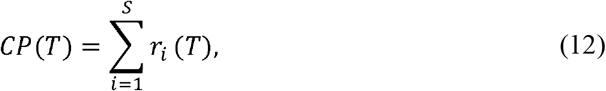

where *r*_*i*_(*T*) is the performance curve of species *i* □. For a set of temperatures *T*, the variability of community performance was quantified as the coefficient of variation:

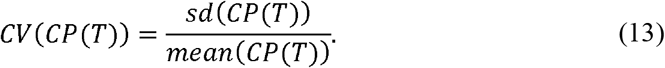

When *T* was the set of actually experienced temperatures, *CV*(*CP*_*actual*_) was the actual CP variability. When the environmental sequence was assumed unknown, we evaluated the CP over a uniform distribution spanning the environmental range; this yielded null CP variability (*CV*(*CP*_*null*_)), which requires no information about the realised frequency of environmental conditions experienced.

In the consumer–resource model, raw growth rate is not expected to map linearly onto realised abundance, because species with positive growth can persist but their realised abundance is determined by resource supply, resource depletion, mortality, and competition (Gonzalez & Feo 2007; Lehman & Tilman 2000). We therefore used a viability transformation (Supplementary Information 2, Section 2.5) that maps intrinsic growth onto whether a species can persist at a given temperature:

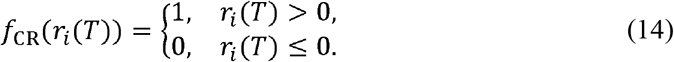

The consumer–resource viability CP was then

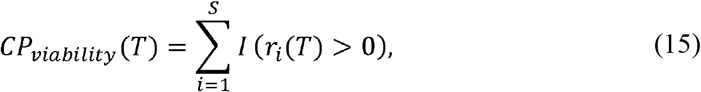

where *I* is an indicator function. This *CP*_*viability*_ is the number of species capable of positive growth at each temperature in the absence of interactions. Its coefficient of variation quantifies variation in the viable species pool across the temperatures over which it is evaluated (e.g. through time for *CP*_*actual*_, or across the specified temperature distribution for *CP*_*null*_).

### Community stability analyses of simulations

For each simulation, community variability (i.e. the inverse of stability) was the coefficient of variation of total abundance through time (*CV*(*N*_tot_)). We compared three predictor sets: actual CP variability (*CV*(*CP*_*actual*_)), null CP variability (*CV*(*CP*_*null*_)), and trait summaries. The trait model included mean thermal mismatch (i.e. the difference between the mean thermal optimum and mean temperature) 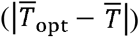, the spread of thermal optima (*sd*(*T*_opt_)), and their interaction:

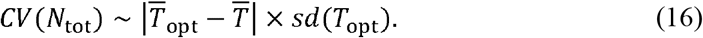

Linear models were fitted separately within each simulation run, interaction-strength or resource-sharing treatment, and temperature-series replicate. Grouped R^2^ therefore measured how well each predictor explained instability across the remaining trait and richness combinations within a common interaction and environmental context.

We also fitted random forests separately to each of the two Lotka–Volterra and two consumer–resource runs, with actual and null CP variability, trait summaries, richness, and interaction strength or resource sharing as predictors, using the ranger R package (Wright & Ziegler 2017). Diagnostics included random-subset robustness (size = 1000 communities), scaled permutation importance, and out-of-bag prediction accuracy.

We compared synchrony predicted from species thermal performance curves with synchrony in abundance. For a community of S species, we used the Loreau-de Mazancourt synchrony metric (Loreau & de Mazancourt 2008):

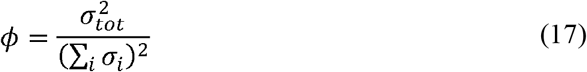

where σ^2^ □ □ □ is the temporal variance of total community abundance and σ□ is the temporal standard deviation of species i; the sum is taken across all S species. Values range from 0 (perfect asynchrony) to 1 (perfect synchrony). We applied the metric both to simulated species abundance and to species performance.

### Empirical ciliate microcosm experiment

We designed a ciliate microcosm experiment to test whether CP variability predicts the temporal stability of total community biomass under fluctuating temperature. We used five aquatic ciliates: *Colpidium striatum, Dexiostoma campylum, Paramecium caudatum, Tetrahymena thermophila*, and *Loxocephalus* sp. Stock cultures were maintained at 15 °C in bacterised alfalfa medium (0.55 g L□^1^) supplemented with 0.5 mL L□^1^ 10× Chalkley’s medium and 1.5 g L□^1^ sodium chloride. Resource bacteria (*Bacillus subtilis, Serratia fonticola*, and *Brevibacillus brevis*) were cultured separately in DMSZ-1 medium at 37 °C for 24 h and combined after three days in filtered alfalfa medium.

First, we measured each species’ temperature response in an independent monoculture experiment. Species were grown at seven constant temperatures (15, 18, 21, 24, 27, 30, and 33 °C) for 15 days, during which the species were sampled 10 times. Growth rate was estimated as the slope of log10 population density during the early growth window for each species-by-temperature combination. This window was chosen separately by species and temperature to capture initial growth before later density-dependent or declining phases. Species-specific thermal performance curves were then fitted to the temperature-specific growth-rate estimates with generalised additive models in the mgcv R package (Wood 2022) (Supplementary Information 3, Figure S1).

The community experiment involved 3 richness levels (2, 3, and 4 species), three levels of CP variability, four temperature treatment, and three replicate for each treatment combination. Temperature treatments consisted of four fluctuating time series. We calculated *CP*_*actual*_ for every possible 2-, 3-, and 4-species community. For each richness level and each of four temperature sequences, we selected the compositions with the lowest *CV*(*CP*_*actual*_) , the highest *CV*(*CP*_*actual*_) , and the value closest to the mean (Supplementary Information 3, Figure S2 and S3). Composition could therefore differ among temperature sequences because actual CP variability depends jointly on species response curves and the environmental sequence. A fully factorial design with three richness levels, three CP-variability classes, four temperature sequences, and three replicate microcosms produced 108 experimental units. Each microcosm contained 25 mL of medium and was maintained in a programmable incubator (IPP260plus, Memmert, Germany).

The four temperature sequences were drawn from a normal distribution with mean 22.5 °C and standard deviation 3.5 °C and had 1/*f* temporal autocorrelation with exponent γ = 0.8, matching the Lotka–Volterra simulations (Supplementary Information 3, Figure S2). Communities acclimated for four days at 22.5 °C before the fluctuating treatments began.

The community experiment ran for 37 days and included 27 sampling days. After excluding the initial four-day constant-temperature period, the analysis contained 25 sampling days. We sampled ciliate densities on each weekday for five weeks using video microscopy. Three microcosm-day observations with sampling errors were removed. For sampling, we transferred 65 μL subsamples to counting chambers and recorded 5-s videos, which were processed with BEMOVI in R (Pennekamp *et al*. 2015, 2017). After sampling, microcosms were replenished with fresh alfalfa medium to maintain volume. Once per week, 75 individuals of each species present in a microcosm were reintroduced to permit recolonisation after local extinction (analogous to immigration terms in the Lotka-Volterra simulations) (Limberger *et al*. 2025).

### Community stability analyses of the experiment

Community biomass was the sum of species biovolumes multiplied by the density of water (1 g cm□^3^). Because biomass declined across species during the experiment (Supplementary Information 3, Figure S4), probably owing to resource depletion or waste accumulation, we removed long-term trends before estimating temporal instability (Isbell *et al*. 2026).

Species-level biomass was detrended with generalised additive models fitted separately to each species-by-composition combination (Fewster *et al*. 2000; Simpson 2018). We summed detrended species deviations within each microcosm and day and quantified empirical variability as the standard deviation of detrended total community biomass:

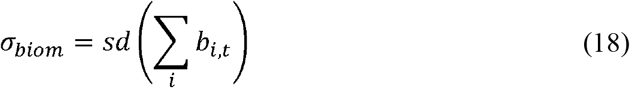

where *b*□, □ is the detrended biomass of species *i* at time t.

Because detrending centres deviations around the fitted long-term trend, we used standard deviation rather than the coefficient of variation since the mean of the detrended series approaches zero which can cause the coefficient of variation to be unstable.

Linear models related log10-transformed community biomass variability (σ_*biom*_) to *CV*(*CP*_*actual*_), *CV*(*CP*_*null*_), 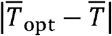, and *sd*(*T*_opt_). CP predictors were log10-transformed to improve normality of residuals; trait-summary predictors were analysed on their original scales. We fitted single-predictor models for each predictor. The main richness-aware model included log10-transformed *CV*(*CP*_*actual*_), richness, temperature time series, and the interaction between *CV*(*CP*_*actual*_) and richness. We included the interaction term considering the results of a previous study where the effect of species’ responses was dependent on the number of species in a community (Polazzo *et al*. 2025). Lastly, we used structural equation modelling (SEM) to investigate hypothesized pathways from *CV*(*CP*_*actual*_) to (*σ*_*biom*_) through synchrony and population variability (Supplementary Information 3, section 2.3.4).

## Results

### Simulations

We quantified explanatory power of the relationship between community abundance variability (*CV*(*N*_tot_)) and various predictors across communities using separate linear models. In the Lotka–Volterra simulations, actual CP variability (*CV*(*CP*_*actual*_)) had a mean R^2^-value of 0.87 and null CP variability (*CV*(*CP*_*null*_)) had a mean R^2^-value of 0.84 (Fig. 2a). R^2^ declined across the interaction-strength gradient, from 0.99 in non-interacting communities to 0.74 in communities with the highest interaction strength for *CV*(*CP*_*actual*_); and from 0.93 to 0.73 for (*CV*(*CP*_*null*_)). The trait-summary model had a mean R^2^ of 0.12 (Fig. 2a) and remained constant across the interaction strength gradient (highest 0.15, lowest 0.13, Fig. 2a).

**Figure 2.**
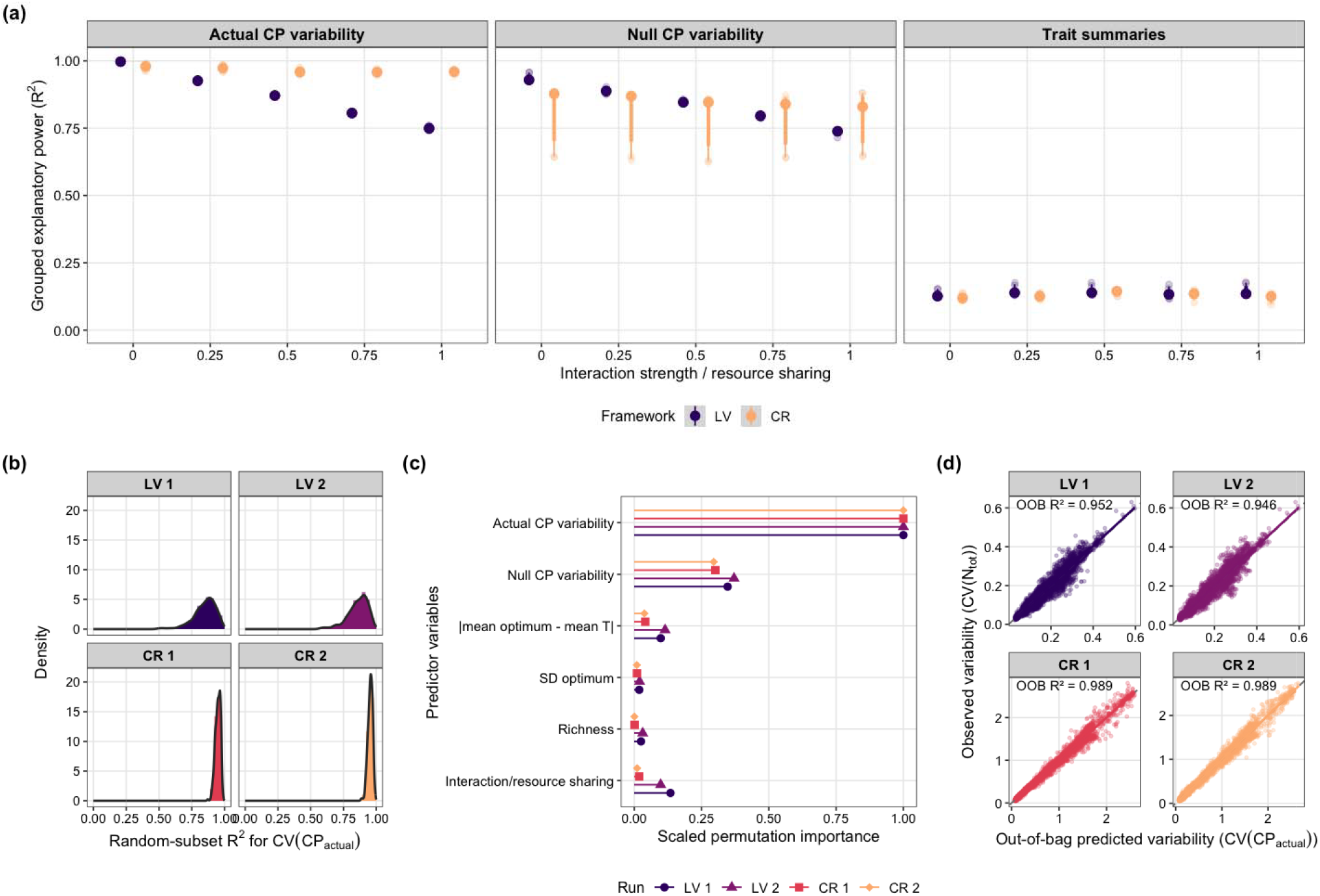
Performance of CP-based predictions in four simulation runs. (a) Explanatory power of actual community-performance variability, null community-performance variability, and trait summaries for temporal variability in total community abundance. Separate linear models were fitted within each simulation run, interaction-strength or resource-sharing treatment, and temperature time series. Small points are the resulting grouped R^2^ values; large points and horizontal bars show the median and 50% and 80% quantile intervals, and shaded half-eye distributions show their density. For the Lotka–Volterra (LV) runs, the horizontal axis is nominal interspecific interaction strength; for the consumer–resource (CR) runs, it is shared-resource use, from entirely private resources (0) to fully shared resource use (1). Actual community performance is evaluated over the experienced temperature sequence, whereas null community performance is evaluated over a uniform distribution spanning the environmental range. LV community performance is summed intrinsic growth rate; CR community performance is the number of species with positive intrinsic growth. The trait-summary model contains absolute mean thermal mismatch, the among-species standard deviation of thermal optima, and their interaction. (b) Robustness of the actual community-performance relationship when each run is randomly partitioned into subsets of 25 communities; histograms and density curves show subset R^2^ values. (c) Permutation importance from 1,000-tree random forests containing actual and null community-performance variability, mean thermal mismatch, thermal-optimum spread, richness, and interaction strength or resource sharing. Importance is scaled within each run so that its largest value equals one. (d) Observed total-abundance variability against out-of-bag random-forest predictions. Points represent communities, solid lines are linear fits, dashed lines show the 1:1 relationship, and labels give out-of-bag R^2^. LV 1 uses asymmetric uniform competition, LV 2 symmetric log-normal competition, CR 1 deterministic resource sharing, and CR 2 stochastic beta-distributed resource sharing. . (c) Scaled random-forest permutation importance. (d) Observed versus out-of-bag predictions.

In the consumer–resource simulations, viability *CV*(*CP*_*actual*_) had a mean R^2^ of 0.96 for models predicting *CV*(*N*_tot_) across the resource-sharing gradient, and the R^2^ did not decline across the interaction strength gradient. Mean R^2^-values were 0.78 for viability (*CV*(*CP*_*null*_)) and 0.12 for the trait-summary model (Fig. 2a).

The relationship between *CV*(*CP*_*actual*_) and *CV*(*N*_tot_) also remained strong across random subsets of the simulation data (Fig. 2b). In a random forest-based analysis of variable importance, *CV*(*CP*_*actual*_) had the highest scaled permutation importance in all four runs after including *CV*(*CP*_*null*_), trait summaries, richness, and interaction strength or resource sharing as predictors (Fig. 2c). Out-of-bag R^2^-values were 0.95 and 0.942 for the two Lotka–Volterra runs and 0.96 for the two consumer–resource runs (Fig. 2d).

In the Lotka–Volterra simulations, the relationship between synchrony predicted from thermal performance and synchrony of abundance declined from R^2^ = 0.99 in the absence of interspecific interactions to R^2^ = 0.066 at the highest interaction strength (Fig. 4 a, and Supplementary Information 2, Figure S10a and b; Table S4). In the consumer–resource model, the corresponding R^2^ declined from 0.67 with entirely private resource use to 0.14 with fully shared resource use (Fig. 4 b, and Supplementary Information 2, Figure S10c and d; Table S4).

**Figure 3.**
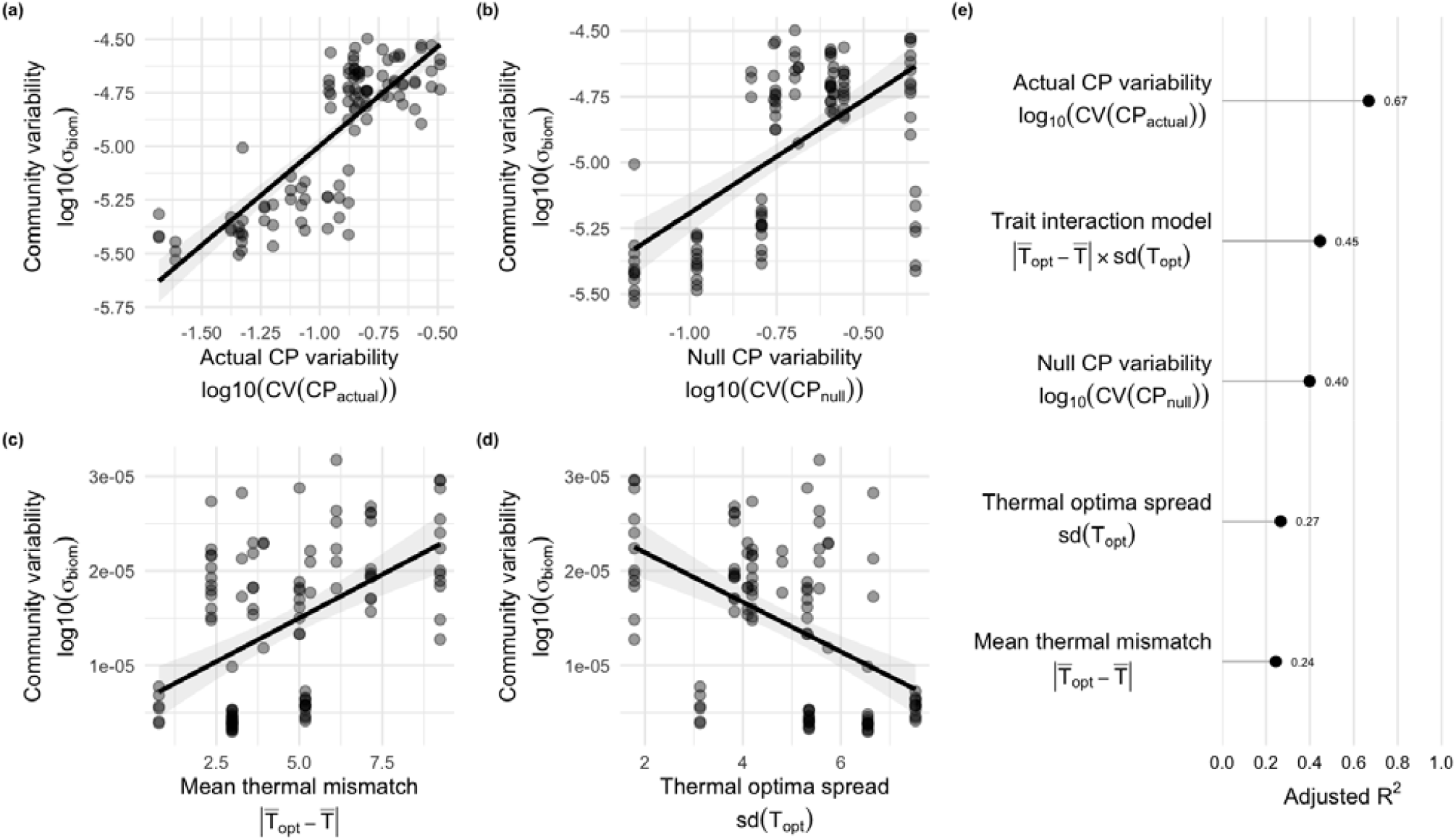
Predictors of detrended community biomass variability. Panels A–D show the relationships between community biomass variability and (A) actual community performance (CP) variability, (B) null CP variability, (C) the absolute difference between mean species thermal optimum and mean experienced temperature, and (D) variation in species thermal optima. Solid lines show fitted linear regressions and shaded areas show 95% confidence intervals. Panel E compares the adjusted values of the four single-predictor models with that of the trait-summary model, which includes mean thermal mismatch, thermal-optimum variation, and their interaction.

**Figure 4.**
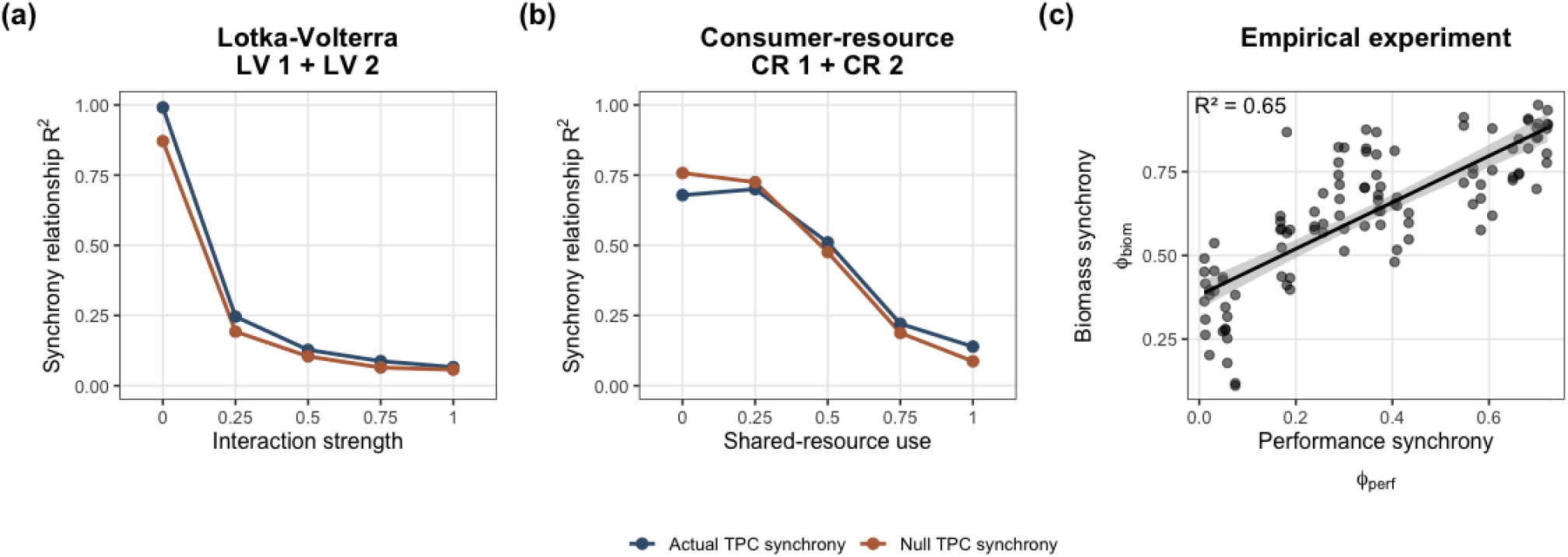
Synchrony links between thermal performance and community dynamics. (a, b) In the Lotka–Volterra and consumer–resource simulations, the relationship between synchrony predicted from species thermal performance and realised abudance synchrony was evaluated separately across interaction-strength or shared-resource-use levels. Lines show the R^2^of realised biomass synchrony explained by performance synchrony calculated from either the actual temperature series or the null temperature distribution. (c) In the empirical ciliate communities, performance synchrony was positively associated with biomass synchrony; points show communities and the line shows the fitted linear relationship with 95% confidence interval.

Derivation of the expected community variability – CP variability relationship for the Lotka-Volterra case explains this high explanatory power despite interaction effects (Supplementary Information 4). When species persist and abundances track the temperature-dependent quasi-equilibrium, variation in CP is expected to approximate a weighted combination of variation in total biomass and weighted species-level abundance variation. The weight *p* depends on the interaction matrix: when the average interaction structure dominates pair-specific heterogeneity, *p* is close to one and should closely track . Calculating *p* for the two Lotka-Volterra experiments showed that realised values were generally high across interaction treatments (median *p* > 0.9), indicating that these simulated communities mostly fall in the regime where CP is expected to remain a strong predictor of biomass stability despite strong interspecific competition.

### Empirical study

*CV* (*CP*_*actual*_)was the strongest single predictor of community biomass variability (*σ*_*biom*_) and explained 67% of the variation (β = 0.92, P < 0.001, Fig. 3a).*CV*(*CP*_*null*_) explained 40% (β = 0.85, P < 0.001, Fig. 3b).Trait summaries in isolation were weaker predictors: greater mismatch between mean community thermal optimum and mean environmental temperature 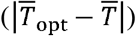 increased variability (*R*^2^=0.25, β = 0.070, P < 0.001, Fig. 3c), whereas greater spread among species’ thermal optima (*sd*(*T*_*opt*_) was stabilizing (*R*^2^=0.27 , β = −0.10, P < 0.001, Fig. 3d). However, a model with an interaction between 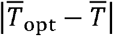 and (*sd*(*T*_*opt*_) explained 45% of the variation in total community biomass variability (Fig. 3e)

Together, *CV* (*CP*_*actual*_) Together, actual , richness, temperature sequence, and the *CV*(*CP*_*actual*_) × richness interaction explained 83% of the variation in community biomass variability (adjusted *R*^2^=0.83, F□, □ □ = 67.11, P < 0.001).*CV*(*CP*_*actual*_) was the most important predictor, while richness, its interaction with CP variability, and temperature sequence each contributed additional explanatory power (all ; *P* < 0.001) Supplementary Information 3, section 2.3.3).

In the experiment, synchrony predicted from species’ thermal performance curves strongly predicted synchrony in their realised biomass fluctuations (= 0.65; Fig. 4c).

## Discussion

Across two independent simulation frameworks and a tailored microcosm experiment, the variability of community performance consistently explained the temporal variability of aggregate community properties. The result held when performance mapped directly onto abundance or did not, when interactions were temperature dependent, and with various assumptions about interspecific interactions and resource sharing patterns. CP variability also outperformed conventional summaries of thermal optima. The central finding is therefore not simply that species responses matter, but that their aggregate shape retains substantial predictive information even after those responses are filtered through ecological interactions.

The synchrony analyses reveal why this result is consequential. Without interspecific interactions, synchrony predicted from thermal performance curves closely matched synchrony in realised abundance. Increasing competition or resource sharing progressively weakened that relationship. Interactions therefore did exactly what interaction-based theory predicts: they reshaped species dynamics and the covariance structure of the community (Gonzalez & Loreau 2009; Hughes & Roughgarden 1998). Yet CP-based predictions of community variability deteriorated only modestly. Aggregate dynamics remained predictable even as the dynamics of their components, and the synchrony among them, became much harder to infer from environmental responses alone.

This apparent decoupling is consistent with classic diversity–stability theory. Community stability depends jointly on population-level variability and synchrony, and interactions can move these components in opposing directions (Ives *et al*. 1999; Lehman & Tilman 2000; Loreau & de Mazancourt 2008; Thibaut & Connolly 2013). Competition may generate compensatory dynamics while simultaneously increasing the variability of individual populations. As a result, changes in covariance can therefore be offset by changes in population variance (Xu *et al*. 2021).

CP variability thus reconciles response-driven and interaction-driven effects on temporal stability. It does not imply that interactions are weak or unimportant: interactions control how environmental forcing shape species dynamics and thus determined the realised synchrony of abundance. Rather, CP variability captures how environmental variation affects the community before interactions redistribute abundance among species. Because the response is aggregated at the same scale as the property being predicted, interaction-driven gains and losses among species can offset, leaving the community-level signal intact. Detailed interaction matrices may therefore be essential for predicting individual populations, but less necessary for forecasting the variability of their aggregate.

The two simulation frameworks also show that CP variability is not a fixed recipe for summing growth rates. It is a way for expressing species responses on the scale that matters for the target community property. Raw growth-based CPs were appropriate in the Lotka–Volterra model, where performance entered population growth directly. In the consumer–resource model, realised abundance depended on resource supply, depletion, and mortality, and a viability transformation was more informative. This contrast is especially informative because the coefficients governing interspecific effects were fixed across temperatures in the Lotka–Volterra model, whereas temperature-dependent resource uptake made effective competition in the consumer–resource model temperature dependent (Duan *et al*. 2025). Environmental variation therefore altered species performance in both models, but could also reshape the interaction strength in the consumer– resource model. The persistence of strong CP-based predictions across this contrast suggests that the framework is robust not only to interactions, but also to interactions whose realised strength changes with the environment. Its weaker null counterpart further shows when knowledge of the realised environmental distribution becomes indispensable when interactions are temperature dependent.

The experiment subjected this framework to a stringent test. Species response curves were estimated independently, communities were selected before the start of the experiment, and no information from the subsequent community experiment was used to adjust the CP predictions. Communities predicted to have more variable aggregate performance were less stable. By predicting community variability before the experiment began, CP variability became a true forecasting tool rather than *a posteriori* explanation. Species richness changed the slope of this relationship, indicating that composition and, consequently, response shape determined how richness influenced stability. Importantly, richness was inherently related to CP variability given the set of species included in this experiment, and we found evidence for a stabilizing pathway of richness through intrinsic performance (Supplementary Information 3, Fig. S8). This finding supports the central hypothesis of response diversity research which proposes intrinsic performance as the underlying mechanism of the diversity-stability relationship (Elmqvist *et al*. 2003; Mori *et al*. 2013). However, the association between richness and CP variability depends on the available species pool and community assemblages and can vary in strength and sign. Thus, shifting the focus to CP variability meaningfully improves predictions.

CP variability outperforms trait summaries because they retain the shape of environmental responses. A mean thermal optimum identifies the centre of a response, and its variance describes dispersion among centres, but neither captures breadth, skew, or overlap. The issue is not merely statistical compression of using trait summaries; it is nonlinear averaging. Because performance curves are nonlinear, performance at the mean environment is not the same as mean performance in a variable environment (Dowd *et al*. 2015; Ruel & Ayres 1999; Ye *et al*. 2019). Jensen’s inequality therefore makes the shape of the response, and the distribution of experienced environments, part of the prediction.

Our tests span competitive and consumer–resource dynamics, a single abiotic driver, and relatively small empirical communities. Trophic interactions, facilitation, multiple covarying drivers, acclimation, and evolutionary change could alter the mapping between performance and abundance. The relevant transformation may also differ among target properties, such as, biomass, productivity, nutrient flux, or organic matter degradation. These need not be considered exceptions to the framework, but rather specifications of it. Use of CP for prediction requires response functions and an aggregation rule matched to the property of interest.

Complete interaction networks remain extraordinarily difficult to estimate in natural ecosystems. Our results suggest that forecasting aggregate temporal stability may often require less information. Environmental variation is first filtered through species response functions, and much of that signal survives the subsequent reorganisation of population dynamics by interactions. Community performance variability provides a scalable way to extract it, offering a route from species-level environmental responses to forecasts of ecosystem stability under environmental change.

## Acknowledgment

The authors want to thank Quqiming Duan for the early analytical work, and the Predictive Ecology group of the University of Zurich for the comments on the draft of the manuscript.

