## Supplementary Information 1 for "Community performance curves predict temporal stability despite interaction effects"

Search strategy, screening criteria, and summary of retained studies

### 1 Methods

#### 1.1 Literature Review

We searched the Web of Science Core Collection on 26 November 2025 for studies linking temporal stability, interspecific interactions, and species-specific environmental responses. The search covered titles, abstracts, and author keywords using the following Topic Search (TS):

```
TS = (("temporal stabilit*" OR "temporal variabilit*")
AND ("interaction*")
AND ("environmental response*" OR "environmental variabilit*"
OR "abiotic factor*" OR "species-environment relationship*"
OR "response diversity" OR "species* performance*"
OR "species respons*" OR "species environment* respons*"))
```

Wildcards captured word stems. We retained peer-reviewed research articles, meta-analyses, and reviews, imposed no date restriction, and excluded conference proceedings, editorials, book chapters, theses, and non-English records.

Records were exported to a reference manager and deduplicated. Two reviewers independently screened titles and abstracts against three criteria: studies had to (i) analyse temporal stability or variability of a community or ecosystem property, (ii) evaluate or discuss interspecific interactions, and (iii) address species-specific environmental responses or related constructs, including response diversity, species-environment relationships, performance curves, or traits linked to abiotic drivers. We excluded studies restricted to spatial stability or environmental gradients without temporal dynamics. Full texts were then screened to confirm eligibility and extract data; disagreements were resolved by discussion.

For each study, we recorded bibliographic information, system and study type, stability metric, representation of interactions, treatment of species-specific responses, and the mechanism judged to prevail when both were considered important. We summarised categorical variables as counts and proportions and visualised associations among study type, species-specific responses, and interactions with Sankey diagrams.

### 2 Results

#### 2.1 Literature Review

The review retained only 12 studies. Ten concluded that species-specific environmental responses were informative for temporal stability and nine identified interactions as important; seven considered both. Of those seven, five judged species-specific responses to be the prevailing mechanism and two judged interactions to prevail.

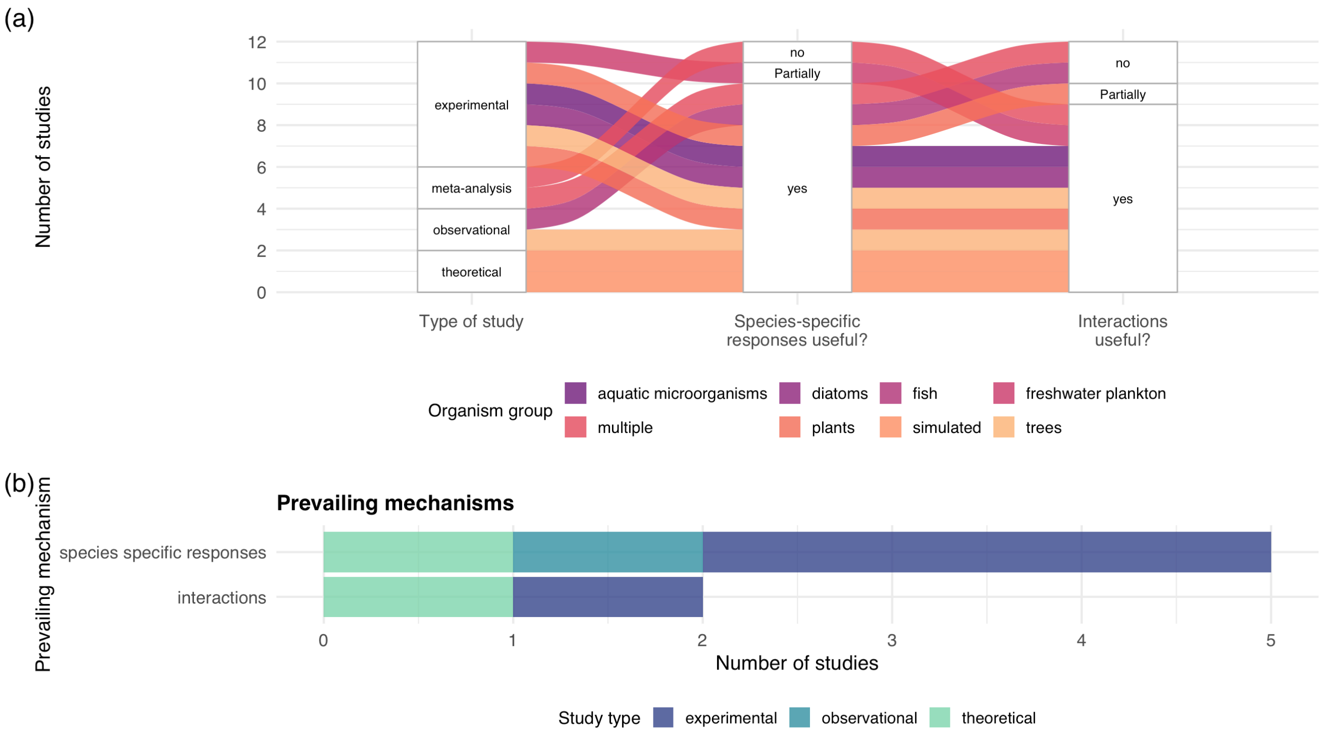


Figure S 1: Systematic review: overview of study characteristics and prevailing mechanisms in the literature. The upper Sankey diagram illustrates the associations between study type, whether species-specific responses were considered important for stability, and whether interactions were considered important for stability. Colours indicate organism groups. The lower bar plot summarizes which mechanism was considered prevailing among studies that concluded both species-specific responses and interactions were important in the Sankey diagram, grouped by study type.
