## Supplementary Information 2 for "Community performance curves predict temporal stability despite interaction effects"

Community performance variability as a predictor of temporal stability

### 1 Overview

This supplementary information presents a reproducible record of the simulations used in the manuscript entitled: Community performance curves predict temporal stability despite interaction effects. It contains the conceptual construction of community performance ($CP$), the simulation experiments, the generalisation of $CP$ through model-specific link functions, and additional simulation analyses.

The simulation section uses two Lotka–Volterra runs and two consumer-resource runs, reported as LV 1, LV 2, CR 1, and CR 2. These runs were designed to test whether temporal variability in community performance predicts temporal variability in total abundance across interaction structures and modelling frameworks.

Figure S 1 shows the raw $CP$ concept. Figure S 2 gives the conceptual construction of viability $CP$. Figure S 3 gives the main simulation result and associated random-forest analysis. Later sections define the model-specific $CP$ construction and compare synchrony predicted from species thermal performance curves with synchrony realised in the simulations.

### 2 Simulation Studies

The simulation analyses use four factorial experiments. LV 1 and LV 2 are Lotka–Volterra simulations in which temperature-dependent performance affects intrinsic growth. CR 1 and CR 2 are consumer-resource simulations in which temperature-dependent uptake and resource sharing generate abundance dynamics through resource-mediated competition. In all simulations, the response variable is temporal variability of total abundance, measured as the coefficient of variation of total abundance.

The main linear models were fitted within each simulation run, interaction-strength or resource-sharing treatment, and temperature series.

#### 2.1 Implementation

Simulation experiments were implemented with the R package `community.simulator`, which provides tools for specifying factorial simulation designs, generating temperature time series, simulating temperature-dependent community dynamics, and calculating community-level summary measures. The workflow used the package functions for reading experiment specifications, creating experiment tables, generating environmental time series, running dynamical simulations, and extracting community measures. The package documentation provides the relevant implementation details in the articles on YAML experiment templates, getting started with a small experiment, and the model-specific walkthroughs for continuous-time Lotka–Volterra and consumer-resource simulations. Function-level documentation is provided for `run_experiment()`, `simulate_dynamics_from_spec()`, `get_community_measures_from_spec()`, `simulator_lv_continuous()`, and `simulator_consumer_resource_continuous()`.

| experiment | framework | cases | treatment_levels | temperature_series | richness | optimum_means | optimu |
|---|---|---|---|---|---|---|---|
| CR1 | Consumer-resource | 7290 | 5 | 3 | 4, 8, 12 | 16, 20, 24 | 0, 2, 4 |
| CR2 | Consumer-resource | 7290 | 5 | 3 | 4, 8, 12 | 16, 20, 24 | 0, 2, 4 |
| LV1 | Lotka-Volterra | 7290 | 5 | 3 | 4, 8, 12 | 16, 20, 24 | 0, 2, 4 |
| LV2 | Lotka-Volterra | 7290 | 5 | 3 | 4, 8, 12 | 16, 20, 24 | 0, 2, 4 |

Table S 1: Publication simulation data included in the supplementary report.

#### 2.2 Conceptual Construction of $CP$

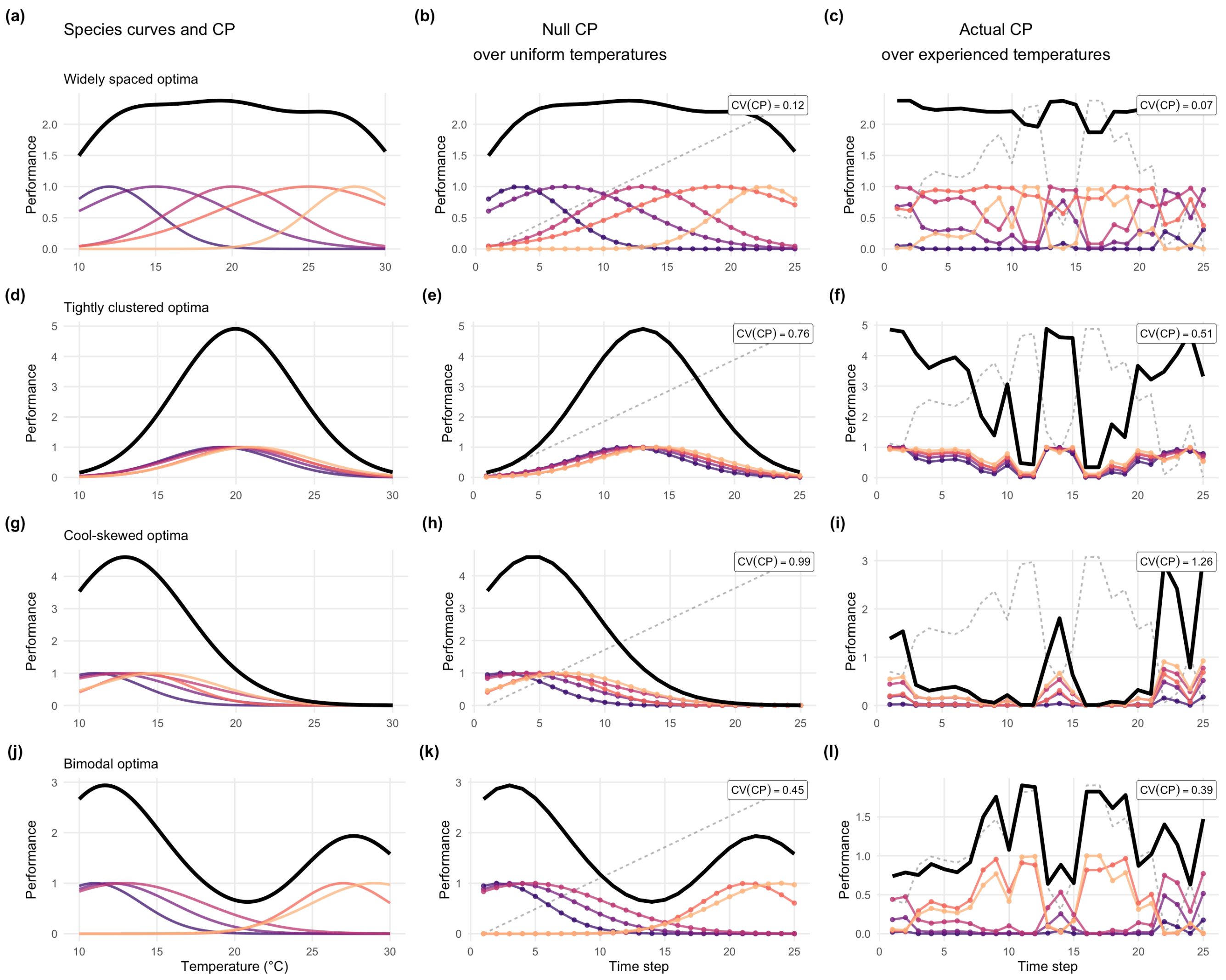

Figure S 1: Conceptual construction of raw community performance ($CP$) using four idealised communities with widely spaced, tightly clustered, cool-skewed, or bimodal species optima. The first column shows species-level performance curves and the summed $CP$ curve. The second column shows null $CP$ time series under uniformly sampled temperatures. The third column shows actual $CP$ time series under an experienced temperature sequence. Dashed grey lines in the time-series panels show the temperature sequence rescaled to the plotting height.

##### 2.2.1 Viability $CP$

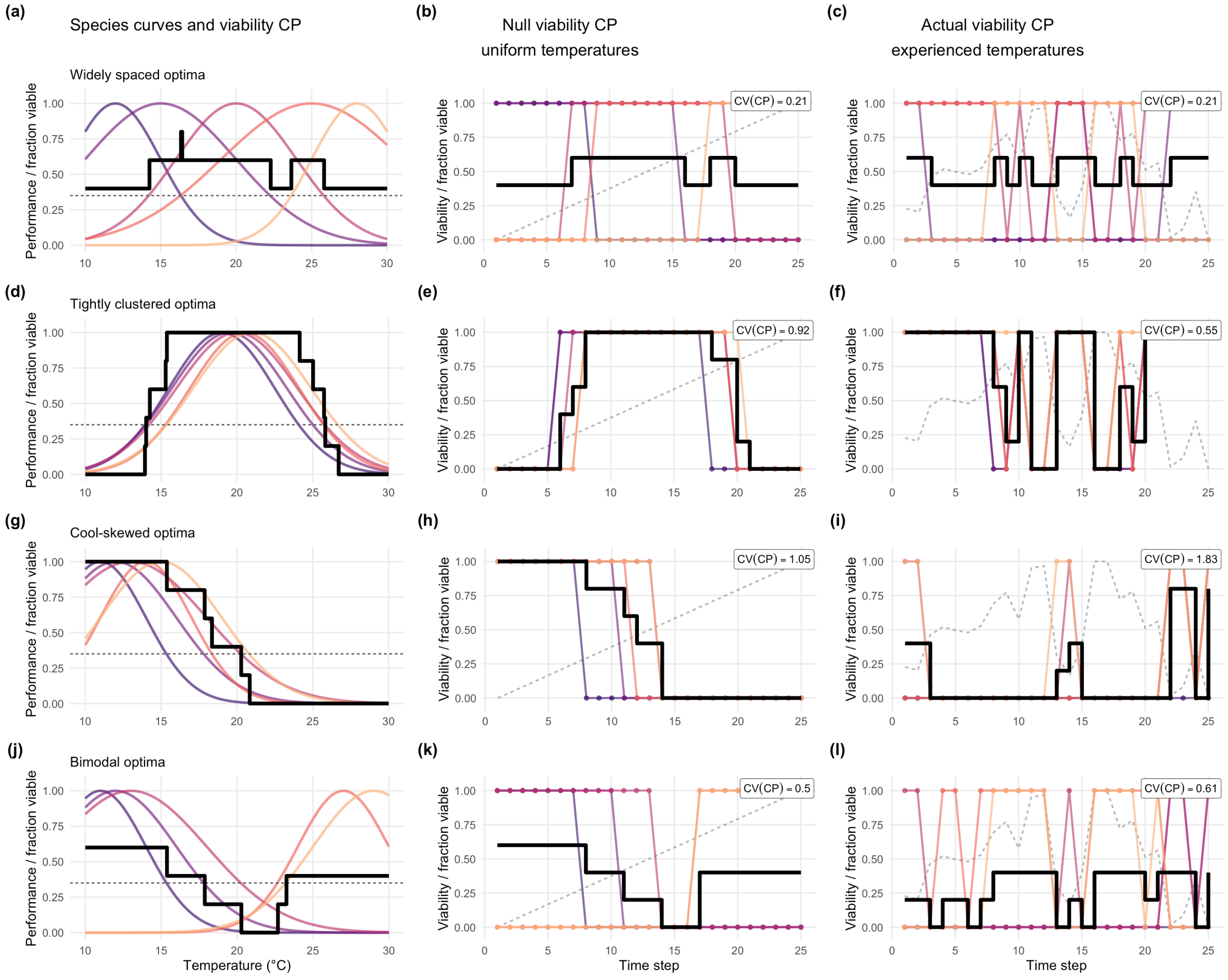


Figure S 2: Conceptual construction of viability $CP$. Species performance curves are first transformed into binary viability using the dashed threshold. Community viability $CP$ is the fraction of species above that threshold at each temperature or time step. Dashed grey traces in the time-series panels show the temperature sequence rescaled to the panel height, and black curves show community-level viability $CP$.

#### 2.3 Simulation Results

Variability of $CP$ predicted community variability across both modelling frameworks, but the relevant $CP$ scale differed between models. In the Lotka–Volterra simulations, where temperature-dependent performance enters directly through intrinsic growth, raw $CV(CP)$ explained a large fraction of variation in total-abundance instability. Actual $CP$ variability, $CV(CP_{\text{actual}})$, had a median grouped (R^2) of 0.87, and null $CP$ variability, $CV(CP_{\text{null}})$, had a very similar median grouped (R^2) of 0.85. By contrast, the trait-summary model explained less variation, with a median grouped (R^2) of 0.13.

In the consumer-resource simulations, raw uptake curves are filtered through resource depletion, mortality, and competition for shared resources. Binary viability therefore provides the $CP$ scale most directly related to total abundance. Actual viability $CP$ variability, $CV(CP_{\mathrm{actual}})$, had a median grouped (R^2) of 0.96. Null viability $CP$ variability, $CV(CP_{\mathrm{null}})$, explained less variation ((R^2 = 0.85)), as did the trait-summary model ((R^2 = 0.13)). These results indicate that $CP$ variability is most informative when $CP$ is constructed on a scale that corresponds to the abundance process being analysed.

The random-forest analysis gave a consistent ranking of predictors. Actual $CP$ variability, $CV(CP_{\mathrm{actual}})$, had the highest permutation importance in all four simulation runs. Null $CP$ variability, $CV(CP_{\mathrm{null}})$, had secondary permutation importance in the Lotka–Volterra runs and lower permutation importance in the consumer-resource runs. Trait summaries, richness, and interaction or resource-sharing treatment had lower permutation importance once $CV(CP_{\mathrm{actual}})$ was included.

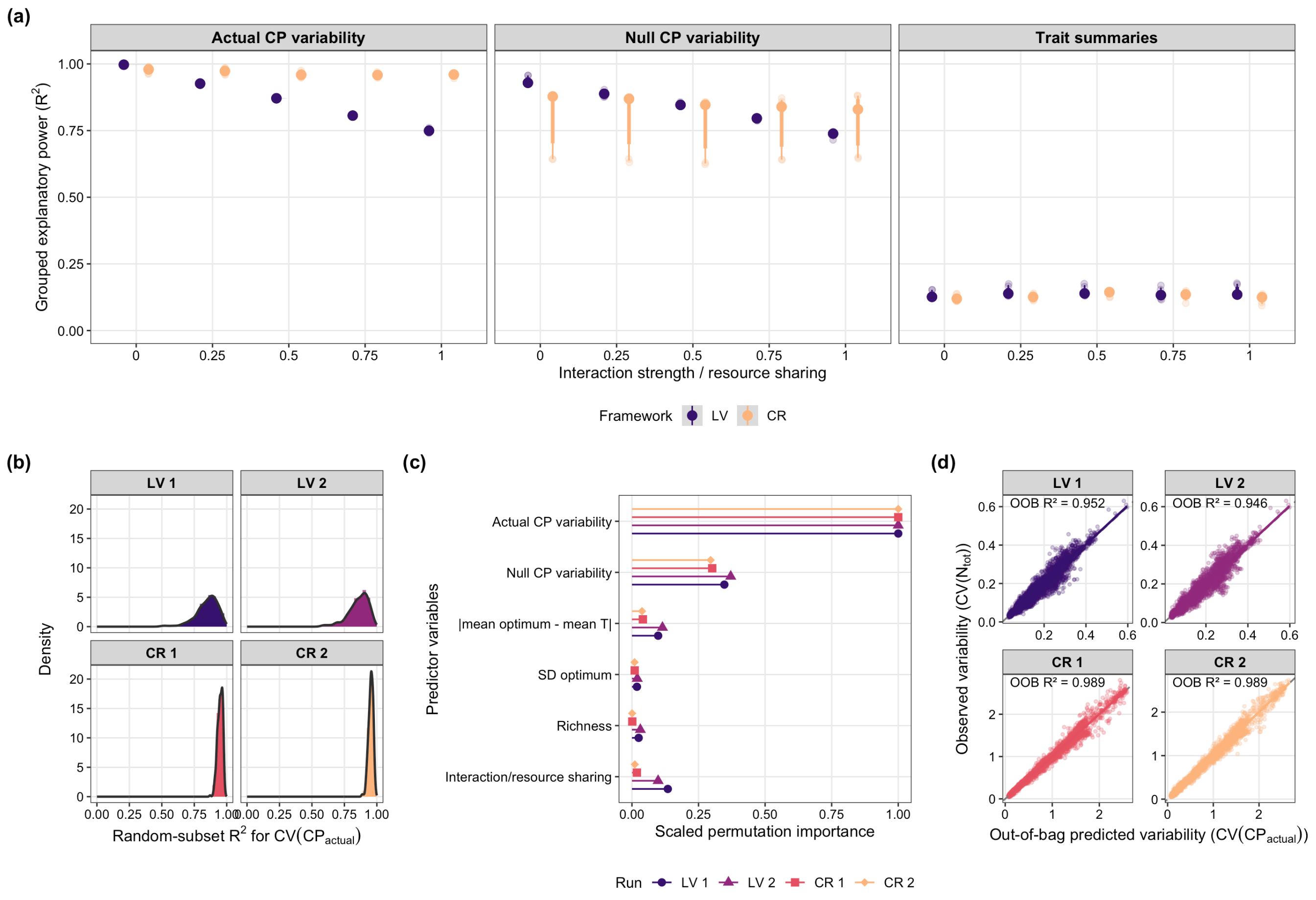


Figure S 3: Explanatory power of community performance metrics across the four simulation experiments. Panel (a) shows linear-model R² values fitted within each run, treatment level, and temperature series; models were not grouped by mean thermal optimum. The x-axis gives nominal interaction strength for LV 1 and LV 2 and resource sharing for CR 1 and CR 2, where 0 indicates no shared resource and 1 indicates fully shared resource use. Facets show actual $CP$ variability ($CV(CP_{\mathrm{actual}})$), null $CP$ variability ($CV(CP_{\mathrm{null}})$), and trait summaries. For LV 1 and LV 2, actual and null $CP$ variability are based on raw community performance. For CR 1 and CR 2, actual and null $CP$ variability are based on binary viability $CP$. Trait summaries are the distance between realised mean optimum and mean environmental temperature, the realised standard deviation of optima, and their interaction. Panel (b) shows random-subset robustness for $CV(CP_{\mathrm{actual}})$. Panel (c) shows random-forest permutation importance for the same predictor set, scaled within each run. Panel (d) compares observed instability with out-of-bag predicted instability.

| framework | experiment_label | grouping | treatment_plot | predictor | groups | mean_r2 |
|---|---|---|---|---|---|---|
| Consumer-resource | CR 1 | Run + temperature series | Pooled | Actual CP variability | 3 | 0.956 |
| Consumer-resource | CR 1 | Run + temperature series | Pooled | Null CP variability | 3 | 0.779 |
| Consumer-resource | CR 1 | Run + temperature series | Pooled | Trait summaries | 3 | 0.123 |
| Consumer-resource | CR 1 | Run + treatment + temperature series | 0 | Actual CP variability | 3 | 0.977 |
| Consumer-resource | CR 1 | Run + treatment + temperature series | 0 | Null CP variability | 3 | 0.802 |
| Consumer-resource | CR 1 | Run + treatment + temperature series | 0 | Trait summaries | 3 | 0.122 |
| Consumer-resource | CR 1 | Run + treatment + temperature series | 0.25 | Actual CP variability | 3 | 0.973 |
| Consumer-resource | CR 1 | Run + treatment + temperature series | 0.25 | Null CP variability | 3 | 0.795 |
| Consumer-resource | CR 1 | Run + treatment + temperature series | 0.25 | Trait summaries | 3 | 0.126 |
| Consumer-resource | CR 1 | Run + treatment + temperature series | 0.5 | Actual CP variability | 3 | 0.957 |
| Consumer-resource | CR 1 | Run + treatment + temperature series | 0.5 | Null CP variability | 3 | 0.772 |
| Consumer-resource | CR 1 | Run + treatment + temperature series | 0.5 | Trait summaries | 3 | 0.141 |
| Consumer-resource | CR 1 | Run + treatment + temperature series | 0.75 | Actual CP variability | 3 | 0.957 |
| Consumer-resource | CR 1 | Run + treatment + temperature series | 0.75 | Null CP variability | 3 | 0.782 |
| Consumer-resource | CR 1 | Run + treatment + temperature series | 0.75 | Trait summaries | 3 | 0.124 |
| Consumer-resource | CR 1 | Run + treatment + temperature series | 1 | Actual CP variability | 3 | 0.962 |
| Consumer-resource | CR 1 | Run + treatment + temperature series | 1 | Null CP variability | 3 | 0.786 |
| Consumer-resource | CR 1 | Run + treatment + temperature series | 1 | Trait summaries | 3 | 0.114 |
| Consumer-resource | CR 2 | Run + temperature series | Pooled | Actual CP variability | 3 | 0.959 |
| Consumer-resource | CR 2 | Run + temperature series | Pooled | Null CP variability | 3 | 0.783 |
| Consumer-resource | CR 2 | Run + temperature series | Pooled | Trait summaries | 3 | 0.128 |

| framework | experiment_label | grouping | treatment_plot | predictor | groups | mean_r2 |
|---|---|---|---|---|---|---|
| Consumer-resource | CR 2 | Run + treatment + temperature series | 0 | Actual CP variability | 3 | 0.977 |
| Consumer-resource | CR 2 | Run + treatment + temperature series | 0 | Null CP variability | 3 | 0.801 |
| Consumer-resource | CR 2 | Run + treatment + temperature series | 0 | Trait summaries | 3 | 0.122 |
| Consumer-resource | CR 2 | Run + treatment + temperature series | 0.25 | Actual CP variability | 3 | 0.972 |
| Consumer-resource | CR 2 | Run + treatment + temperature series | 0.25 | Null CP variability | 3 | 0.792 |
| Consumer-resource | CR 2 | Run + treatment + temperature series | 0.25 | Trait summaries | 3 | 0.125 |
| Consumer-resource | CR 2 | Run + treatment + temperature series | 0.5 | Actual CP variability | 3 | 0.962 |
| Consumer-resource | CR 2 | Run + treatment + temperature series | 0.5 | Null CP variability | 3 | 0.781 |
| Consumer-resource | CR 2 | Run + treatment + temperature series | 0.5 | Trait summaries | 3 | 0.138 |
| Consumer-resource | CR 2 | Run + treatment + temperature series | 0.75 | Actual CP variability | 3 | 0.960 |
| Consumer-resource | CR 2 | Run + treatment + temperature series | 0.75 | Null CP variability | 3 | 0.782 |
| Consumer-resource | CR 2 | Run + treatment + temperature series | 0.75 | Trait summaries | 3 | 0.139 |
| Consumer-resource | CR 2 | Run + treatment + temperature series | 1 | Actual CP variability | 3 | 0.955 |
| Consumer-resource | CR 2 | Run + treatment + temperature series | 1 | Null CP variability | 3 | 0.786 |
| Consumer-resource | CR 2 | Run + treatment + temperature series | 1 | Trait summaries | 3 | 0.125 |
| Lotka-Volterra | LV 1 | Run + temperature series | Pooled | Actual CP variability | 3 | 0.840 |
| Lotka-Volterra | LV 1 | Run + temperature series | Pooled | Null CP variability | 3 | 0.811 |
| Lotka-Volterra | LV 1 | Run + temperature series | Pooled | Trait summaries | 3 | 0.134 |
| Lotka-Volterra | LV 1 | Run + treatment + temperature series | 0 | Actual CP variability | 3 | 0.997 |
| Lotka-Volterra | LV 1 | Run + treatment + temperature series | 0 | Null CP variability | 3 | 0.938 |
| Lotka-Volterra | LV 1 | Run + treatment + temperature series | 0 | Trait summaries | 3 | 0.134 |

| framework | experiment_label | grouping | treatment_plot | predictor | groups | mean_r2 |
|---|---|---|---|---|---|---|
| Lotka-Volterra | LV 1 | Run + treatment + temperature series | 0.25 | Actual CP variability | 3 | 0.932 |
| Lotka-Volterra | LV 1 | Run + treatment + temperature series | 0.25 | Null CP variability | 3 | 0.893 |
| Lotka-Volterra | LV 1 | Run + treatment + temperature series | 0.25 | Trait summaries | 3 | 0.142 |
| Lotka-Volterra | LV 1 | Run + treatment + temperature series | 0.5 | Actual CP variability | 3 | 0.873 |
| Lotka-Volterra | LV 1 | Run + treatment + temperature series | 0.5 | Null CP variability | 3 | 0.849 |
| Lotka-Volterra | LV 1 | Run + treatment + temperature series | 0.5 | Trait summaries | 3 | 0.142 |
| Lotka-Volterra | LV 1 | Run + treatment + temperature series | 0.75 | Actual CP variability | 3 | 0.809 |
| Lotka-Volterra | LV 1 | Run + treatment + temperature series | 0.75 | Null CP variability | 3 | 0.795 |
| Lotka-Volterra | LV 1 | Run + treatment + temperature series | 0.75 | Trait summaries | 3 | 0.145 |
| Lotka-Volterra | LV 1 | Run + treatment + temperature series | 1 | Actual CP variability | 3 | 0.743 |
| Lotka-Volterra | LV 1 | Run + treatment + temperature series | 1 | Null CP variability | 3 | 0.733 |
| Lotka-Volterra | LV 1 | Run + treatment + temperature series | 1 | Trait summaries | 3 | 0.148 |
| Lotka-Volterra | LV 2 | Run + temperature series | Pooled | Actual CP variability | 3 | 0.850 |
| Lotka-Volterra | LV 2 | Run + temperature series | Pooled | Null CP variability | 3 | 0.820 |
| Lotka-Volterra | LV 2 | Run + temperature series | Pooled | Trait summaries | 3 | 0.137 |
| Lotka-Volterra | LV 2 | Run + treatment + temperature series | 0 | Actual CP variability | 3 | 0.997 |
| Lotka-Volterra | LV 2 | Run + treatment + temperature series | 0 | Null CP variability | 3 | 0.938 |
| Lotka-Volterra | LV 2 | Run + treatment + temperature series | 0 | Trait summaries | 3 | 0.134 |
| Lotka-Volterra | LV 2 | Run + treatment + temperature series | 0.25 | Actual CP variability | 3 | 0.920 |
| Lotka-Volterra | LV 2 | Run + treatment + temperature series | 0.25 | Null CP variability | 3 | 0.881 |
| Lotka-Volterra | LV 2 | Run + treatment + temperature series | 0.25 | Trait summaries | 3 | 0.151 |

| framework | experiment_label | grouping | treatment_plot | predictor | groups | mean_r2 |
|---|---|---|---|---|---|---|
| Lotka-Volterra | LV 2 | Run + treatment + temperature series | 0.5 | Actual CP variability | 3 | 0.871 |
| Lotka-Volterra | LV 2 | Run + treatment + temperature series | 0.5 | Null CP variability | 3 | 0.846 |
| Lotka-Volterra | LV 2 | Run + treatment + temperature series | 0.5 | Trait summaries | 3 | 0.151 |
| Lotka-Volterra | LV 2 | Run + treatment + temperature series | 0.75 | Actual CP variability | 3 | 0.803 |
| Lotka-Volterra | LV 2 | Run + treatment + temperature series | 0.75 | Null CP variability | 3 | 0.794 |
| Lotka-Volterra | LV 2 | Run + treatment + temperature series | 0.75 | Trait summaries | 3 | 0.129 |
| Lotka-Volterra | LV 2 | Run + treatment + temperature series | 1 | Actual CP variability | 3 | 0.757 |
| Lotka-Volterra | LV 2 | Run + treatment + temperature series | 1 | Null CP variability | 3 | 0.738 |
| Lotka-Volterra | LV 2 | Run + treatment + temperature series | 1 | Trait summaries | 3 | 0.147 |

Table S 2: Mean grouped explanatory power by modelling framework, grouping scheme, and predictor. Treatment-level rows come from linear models fitted within each run, interaction-strength or resource-sharing treatment, and temperature series. Pooled-treatment rows show the corresponding grouped $R^2$ values when treatment levels are not used for grouping; these models are fitted within each run and temperature series only.

#### 2.4 Simulation Interpretation

The simulations support two linked conclusions. First, variability of $CP$ predicts temporal community variability across both modelling frameworks, despite the fact that the frameworks differ substantially in how species responses enter abundance dynamics. Second, the appropriate $CP$ scale depends on the model. In the Lotka–Volterra simulations, raw $CV(CP)$ is appropriate because temperature-dependent performance enters directly into intrinsic growth. In the consumer-resource simulations, abundance depends on resource supply, depletion, mortality, and competition for shared resources. Binary viability $CP$ is therefore the more relevant community response for predicting total abundance variability.

The consumer-resource simulations address a limitation of the Lotka–Volterra simulations. In the Lotka–Volterra model, interaction coefficients are not temperature dependent. In the consumer-resource model, uptake rates are temperature dependent and species compete through shared resources, implying temperature-dependent effective interaction strengths. The high predictive power of actual viability $CP$ variability, $CV(CP_{\text{actual}})$, in CR 1 and CR 2 therefore indicates that the main $CP$-stability result is robust to a mechanistic setting in which interaction strengths can vary with temperature.

#### 2.5 Constructing $CP$ on an Appropriate Scale

##### 2.5.1 General Definition

A community performance curve ($CP$) can be defined by first mapping each species' temperature-dependent performance onto the scale of the community variable whose stability is being analysed, and then summing across species.

For stability of total abundance or total biomass, the species-level curve therefore represents expected contribution to abundance or biomass, not just raw physiological performance or intrinsic growth rate.

The general construction is:

1. Compute each species' temperature-dependent performance curve.
2. Transform each species' performance to the scale of the focal community variable.
3. Sum the transformed species curves to obtain the $CP$.
4. Use the temporal variability of that $CP$ to predict the stability of the focal community variable.

In notation:

$$P_i(T) = \text{raw performance of species } i \text{ at temperature } T$$

$$A_i(T) = f_{\text{model}, y}(P_i(T))$$

$$CP_y(T) = \sum_i A_i(T)$$

where $y$ is the community variable whose stability is being predicted, such as total abundance or total biomass. The function $f_{\text{model}, y}$ is a model-specific link from raw species performance to expected contribution to that community variable.

##### 2.5.2 Why The Link Function Matters

When predicting the stability of total abundance or total biomass, the relevant question is not simply whether species have high or low raw performance. The relevant question is how changes in species performance translate into changes in species abundance or biomass.

If raw performance is approximately linearly related to abundance, then summing raw performance curves can be appropriate. But if abundance responds nonlinearly, or if performance mainly determines whether a species can persist at all, then raw performance is not on the scale of the response variable. In that case, the species performance curve is first transformed onto an abundance or biomass scale before constructing the $CP$.

This gives the following definition:

> The $CP$ is a sum of species' expected contributions to the focal community variable, not necessarily a sum of raw species performance curves.

##### 2.5.3 Lotka-Volterra Model

In the Lotka-Volterra model used here, raw temperature-dependent performance is closely linked to expected abundance by the model structure. Specifically, the model links intrinsic growth rate and carrying capacity in a way that makes higher performance correspond approximately linearly to higher expected abundance.

For predicting stability of total abundance, the link function from performance to abundance-scale contribution can be treated as approximately identity:

$$f_{\text{LV}, abundance}(P_i(T)) \approx P_i(T)$$

So the LV abundance-scale $CP$ can be written as:

$$CP_{\text{LV}}(T) = \sum_i P_i(T)$$

Each species contributes its raw performance value to the community performance curve. In the Lotka–Volterra simulations, the scale of the summed curve is therefore close to the scale of the abundance response.

##### 2.5.4 Consumer-Resource Model

In the consumer-resource model, raw species performance is not linearly related to equilibrium abundance. Instead, intrinsic growth rate mainly determines whether a species can persist under the current thermal and resource context.

If growth is negative, the species declines to very low abundance. If growth is positive, the species can grow, but its eventual abundance is then determined by other conditions, including:

- resource supply;
- mortality;
- conversion efficiency;
- half-saturation constants;
- competition with other consumers for shared resources.

For predicting stability of total abundance, the abundance-scale transformation is therefore represented by a viability threshold:

$$f_{\text{CR}, abundance}(P_i(T)) = \begin{cases} 1, & P_i(T) > 0 \\ 0, & P_i(T) \leq 0 \end{cases}$$

The consumer-resource abundance-scale $CP$ is then:

$$CP_{\text{CR viability}}(T) = \sum_i I(P_i(T) > 0)$$

where $I(P_i(T) > 0)$ is 1 when species $i$ has positive growth at temperature $T$, and 0 otherwise.

The $CP$ is the number of viable species through time. Its $CV$ measures temporal variation in the viable species pool under the temperature sequence.

##### 2.5.5 Raw and Viability $CP$

Raw $CV(CP)$ sums species growth-rate or performance curves. This construction is appropriate for the Lotka–Volterra model because raw performance is approximately on the abundance scale. In the consumer-resource model, raw growth rate is not the abundance response. It primarily controls whether a species can persist, while the abundance of persistent species is shaped by resource-mediated interactions.

The viability-transformed $CP$ applies a model-specific link function before summing:

$$\text{raw performance} \rightarrow \text{expected abundance-scale contribution} \rightarrow \text{community performance curve}$$

##### 2.5.6 Proposed General Definition

For any model and any focal community variable, define a $CP$ as:

$$CP_y(T) = \sum_i f_{\text{model},\, y}(P_i(T))$$

where:

- $P_i(T)$ is the raw species performance curve;
- $y$ is the community variable whose stability is being predicted;
- $f_{\text{model},\, y}$ maps raw species performance to expected contribution to $y$;
- the $CP$ is formed only after this transformation.

For the current models and response variable:

| Model | Focal stability variable | Link from species performance to contribution |
|---|---|---|
| Lotka-Volterra | Total abundance | Approximately identity |
| Consumer-resource | Total abundance | Binary viability, I(P_i(T) > 0) |

Table S 3: Model-specific $CP$ constructions for total-abundance stability.

This definition keeps the $CP$ concept general while allowing the species-level performance-to-abundance link to differ among models.

##### 2.5.7 Model-Link Cases

The figure panels below apply this construction to the simulation outputs. Total-abundance variability is shown as $CV_{\text{total abundance}}$. The x-axis is $CV(CP)$ constructed under each model-link combination.

For the Lotka-Volterra model, the identity-link $CP$ comes directly from the community-measures output. The binary-link $CP$ uses the viability $CP$ calculated for the same runs. For the consumer-resource model, both identity-link and binary-link $CV(CP)$ values come from the community-measures output.

###### 2.5.7.1 Lotka-Volterra, Identity Link

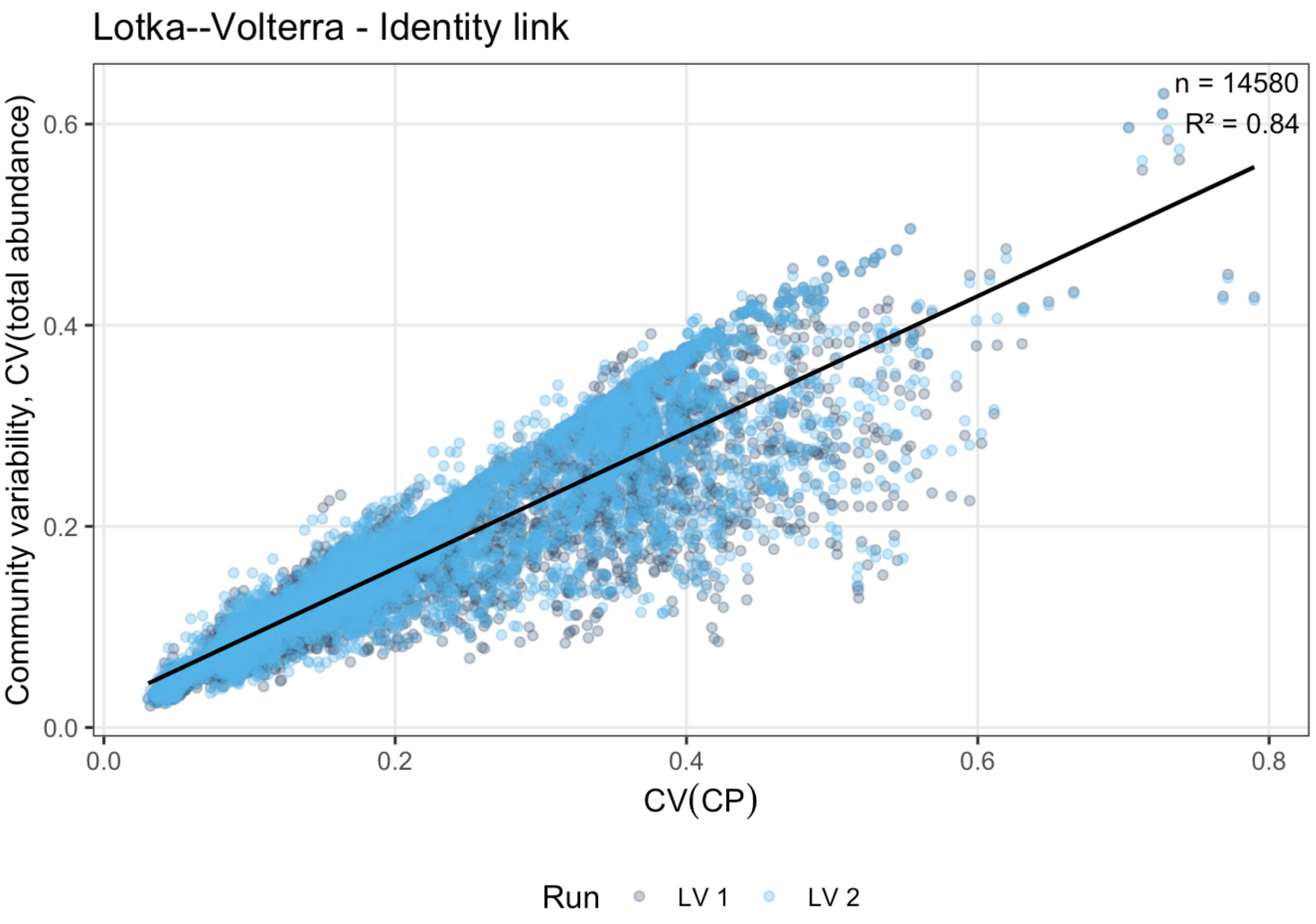


Figure S 4: Lotka-Volterra identity-link $CP$. Points show simulation cases, the x-axis shows $CV(CP)$ based on raw community performance, and the y-axis shows total-abundance variability as $CV(\text{total abundance})$. The identity link is the model-consistent $CP$ construction for the Lotka-Volterra simulations.

##### 2.5.7.2 Lotka-Volterra, Binary Link

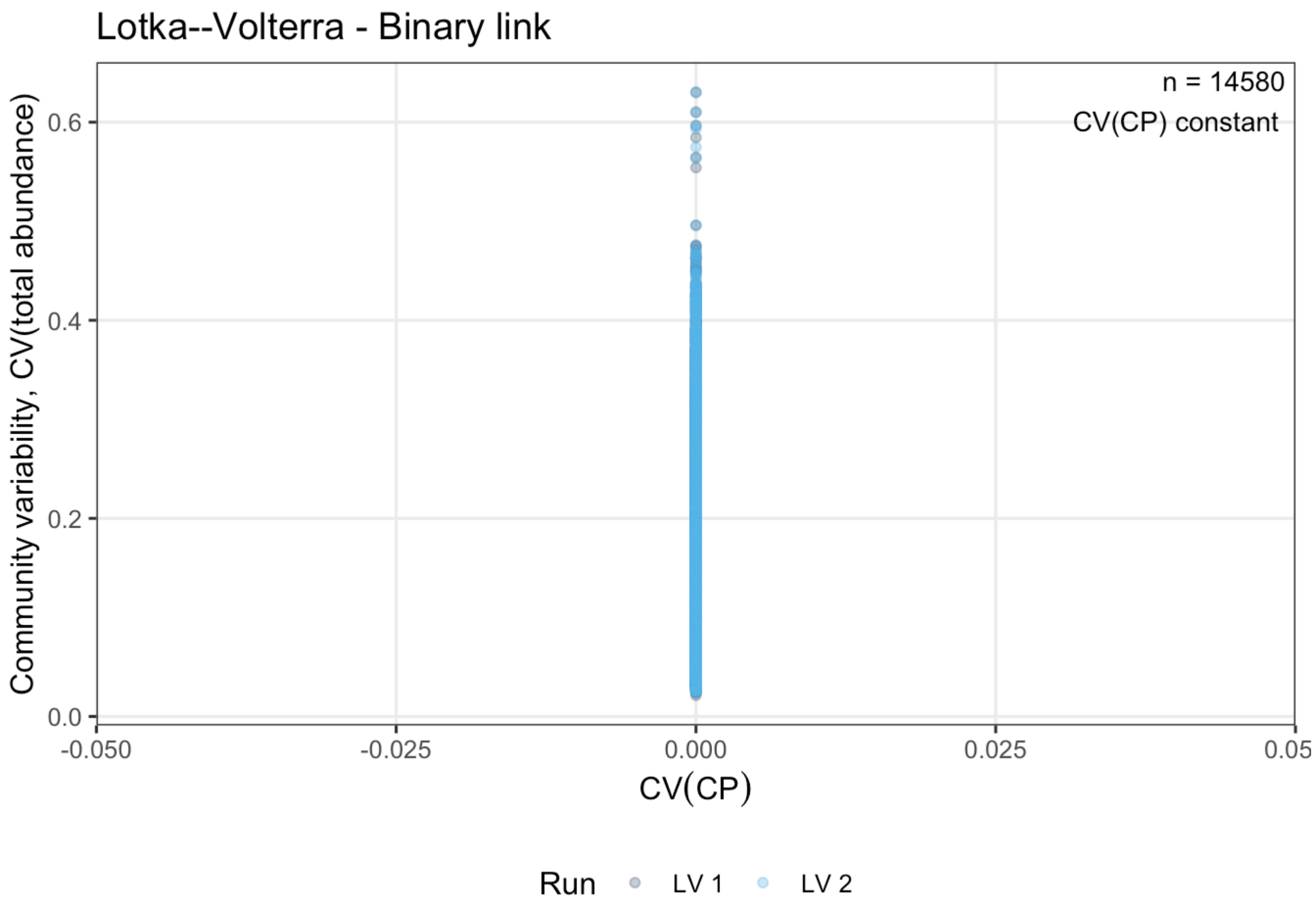


Figure S 5: Lotka-Volterra binary-link $CP$. Species performance is transformed into binary viability before constructing $CP$. This comparison evaluates the effect of reducing the Lotka-Volterra abundance response to a viability count rather than using the raw performance scale.

The binary-link $CP$ evaluates the effect of reducing the Lotka–Volterra response to a viability count rather than using the raw performance scale that enters the model dynamics directly.

###### 2.5.7.3 Consumer-Resource, Identity Link

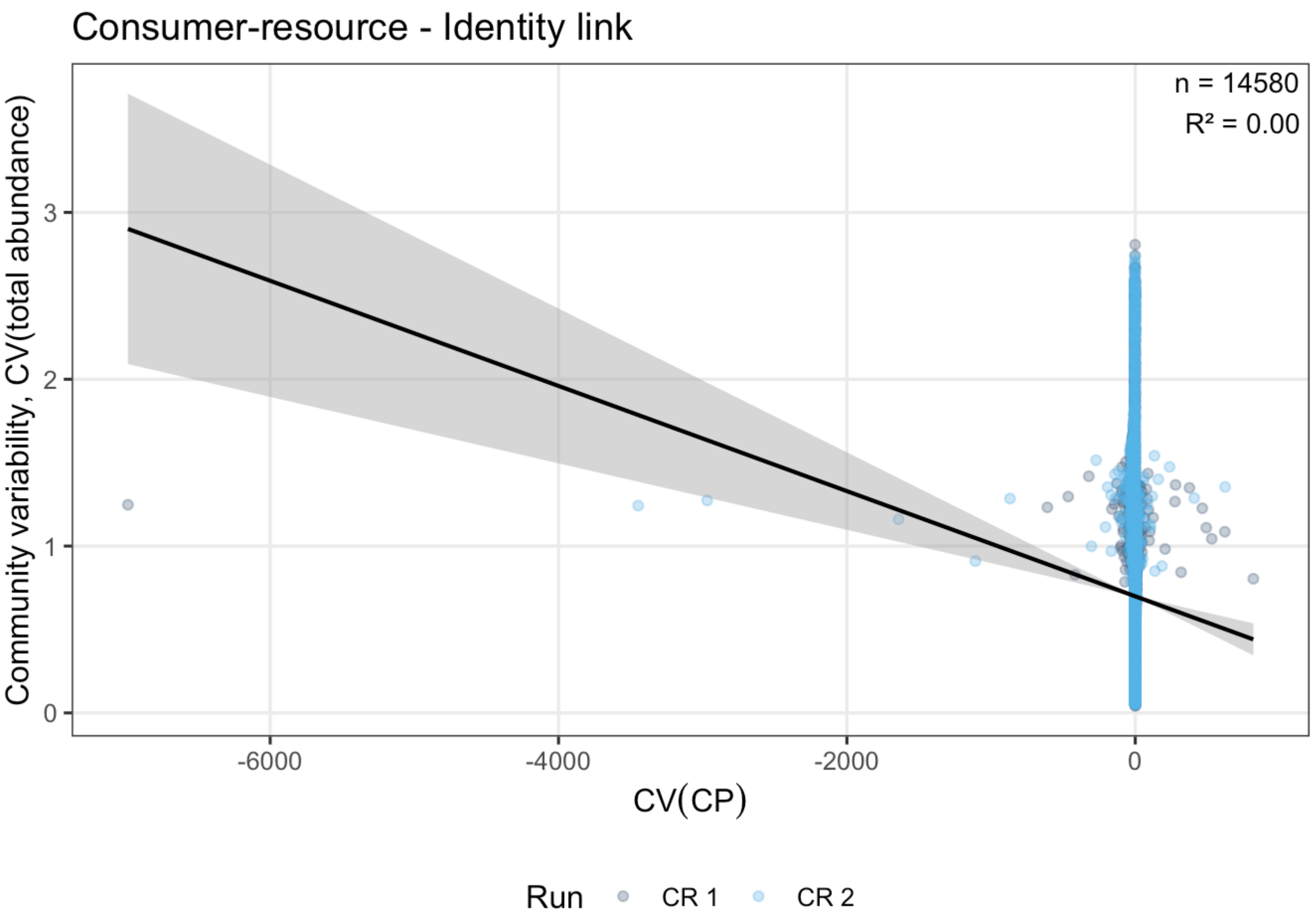


Figure S 6: Consumer-resource identity-link $CP$. Raw performance values are summed directly for the consumer-resource simulations, without first mapping performance onto an abundance-relevant viability scale.

The identity-link $CP$ in the consumer-resource model sums raw uptake or raw performance directly rather than first mapping performance onto an abundance-relevant viability scale.

The same relationship with $CV(CP)$ values above 50 excluded is:

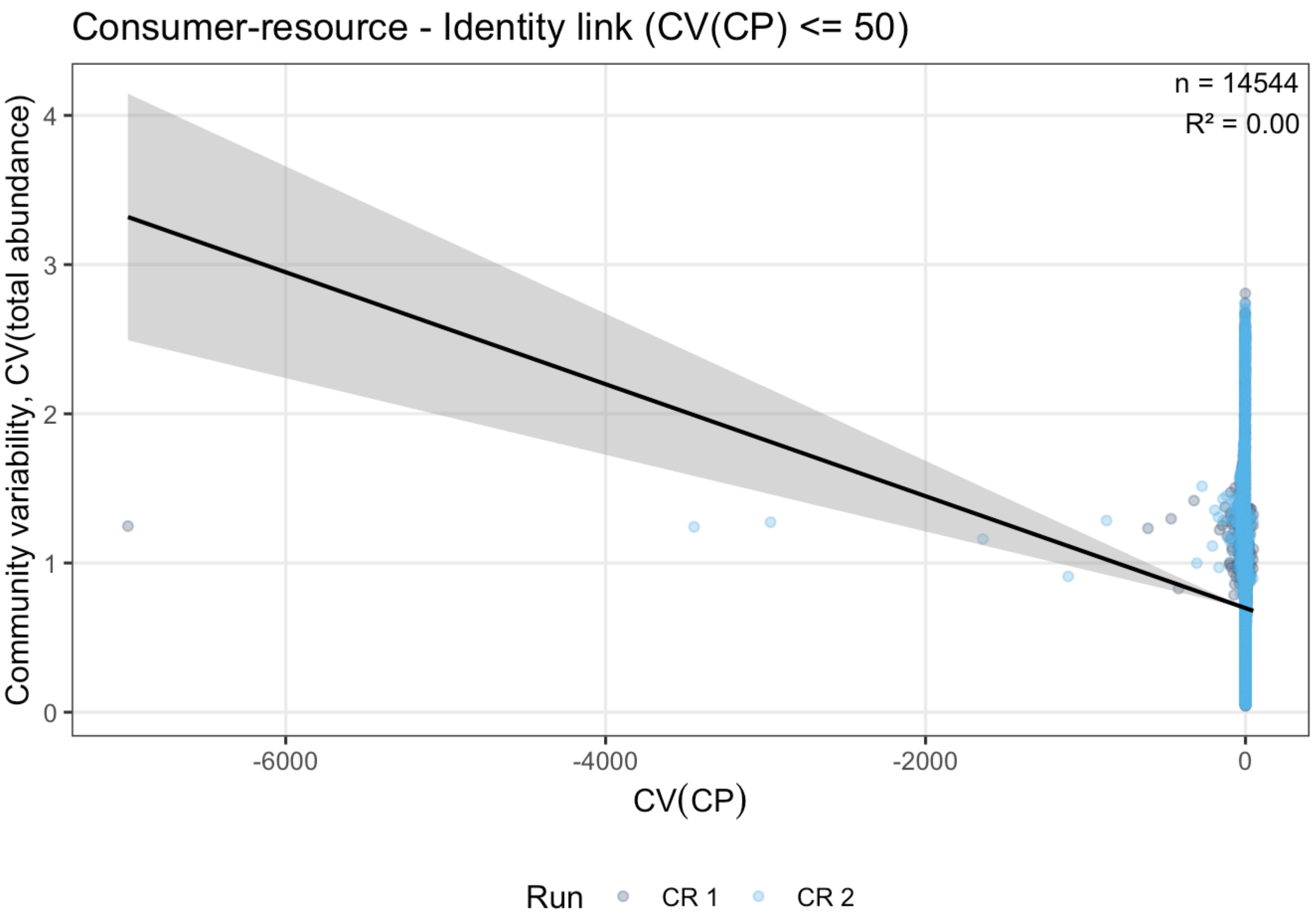


Figure S 7: Consumer-resource identity-link $CP$ after excluding extreme raw $CV(CP)$ values. The filter removes cases with $CV(CP)$ above the stated threshold to show the relationship over the visually interpretable range of the raw identity-link metric.

##### 2.5.7.4 Consumer-Resource, Binary Link

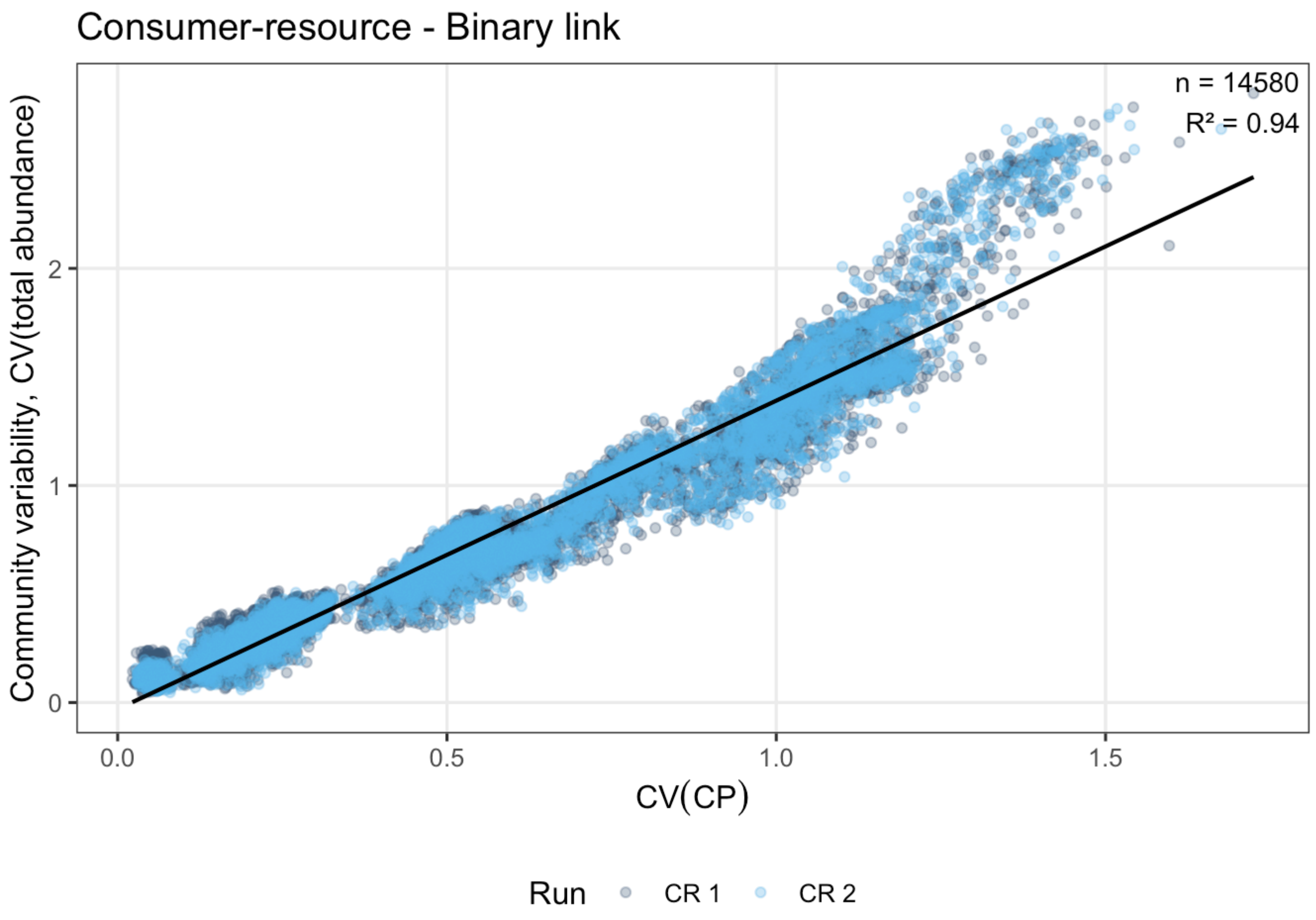


Figure S 8: Consumer-resource binary-link $CP$. Species performance is transformed into binary viability before constructing $CP$, matching the interpretation that total-abundance stability in the consumer-resource simulations is linked to the stability of the viable species pool.

The identity link was most closely aligned with the Lotka–Volterra model, whereas the binary viability link was most closely aligned with the consumer-resource model. The alternative link functions show how predictive performance changes when $CP$ is constructed on a scale less directly tied to the abundance variable whose stability is being predicted.

##### 2.5.8 Practical Interpretation

When the focal response is total abundance or biomass stability, $CP$ addresses the following question:

> How much total abundance or biomass would the community be expected to support at each temperature, given the species that can contribute under that model?

In the LV model, species' raw performance values already approximate their expected abundance contributions. In the consumer-resource model, the first-order abundance contribution is closer to whether each species is viable at all.

The predictive performance of viability $CP$ variability, $CV(CP)$, in the consumer-resource experiments supports the interpretation that $CP$ is most informative when constructed on the scale of the community variable whose stability is being predicted.

##### 2.5.9 Summary of Raw and Viability $CP$

[Figure S 9](#) summarises the same link-function comparison as explanatory power for actual-temperature $CP$ constructions, using the total-abundance instability response used in the main simulation analyses.

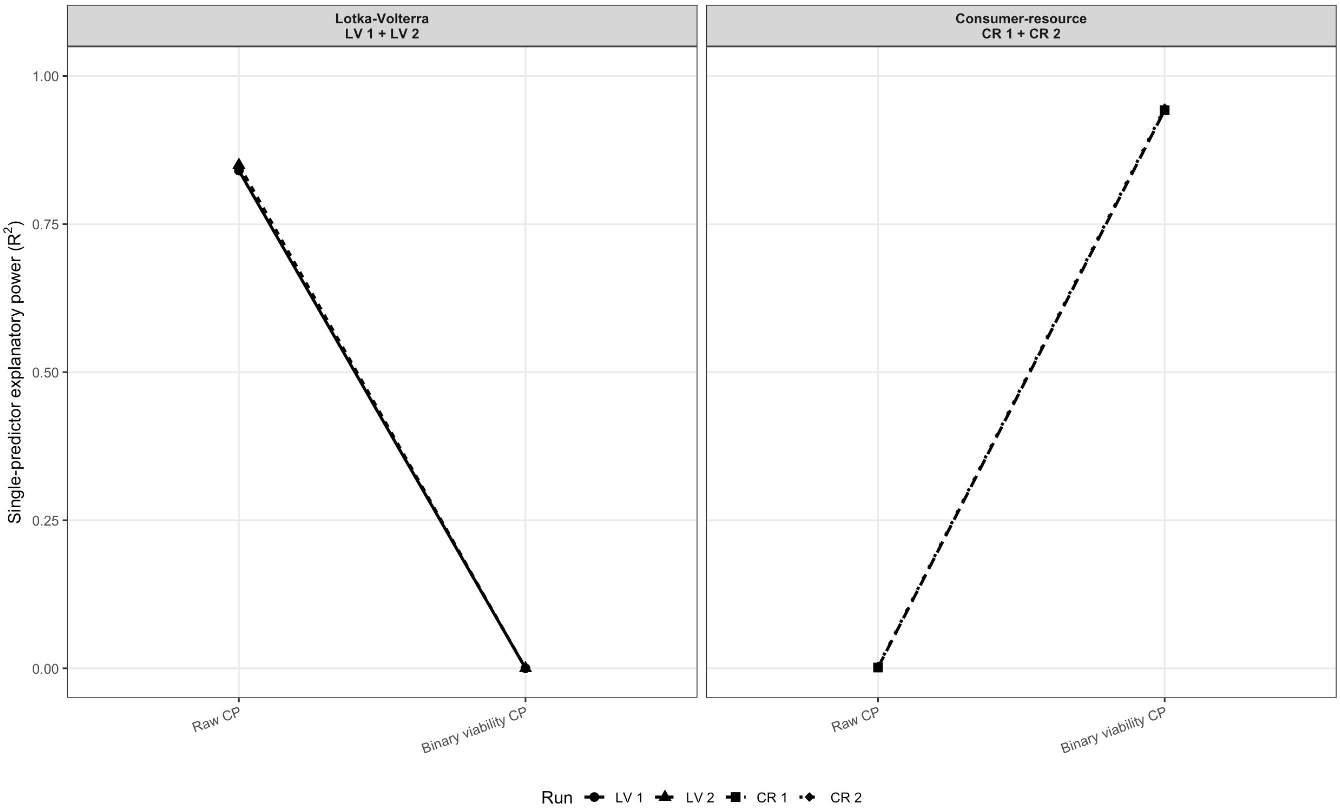


Figure S 9: Comparison of raw and binary $CP$ constructions for the Lotka-Volterra and consumer-resource simulations. Each point is the explanatory power of a single-predictor model relating total-abundance instability to one actual-temperature $CP$ variability metric, fitted separately for each run. Raw $CP$ sums expected performance or growth-rate values directly. Binary viability $CP$ counts species with positive expected net growth.

The comparison supports the link-function interpretation. Actual binary and raw $CP$ metrics differ in their explanatory power across modelling frameworks. In the consumer-resource simulations, the binary viability scale corresponds more closely to the abundance dynamics generated by resource-mediated competition.

#### 2.6 Additional Simulation Analyses

The supplementary analysis tests whether species thermal performance curves predict the synchrony realised in the dynamical simulations, and whether this relationship depends on interaction strength or resource sharing.

##### 2.6.1 Predicted Versus Realised Synchrony

Synchrony was calculated with the Loreau–de Mazancourt metric. For a set of species-level time series $X_i(t)$, synchrony was

$$\phi = \frac{\sigma^2\left(\sum_i X_i(t)\right)}{\left(\sum_i \sigma(X_i(t))\right)^2}.$$

For realised synchrony, $X_i(t)$ was the simulated abundance of species $i$. For TPC-predicted synchrony, $X_i(t)$ was the species performance expected under either the realised temperature sequence or the null temperature distribution. Values near 0 indicate asynchronous species-level fluctuations, whereas values near 1 indicate synchronous fluctuations.

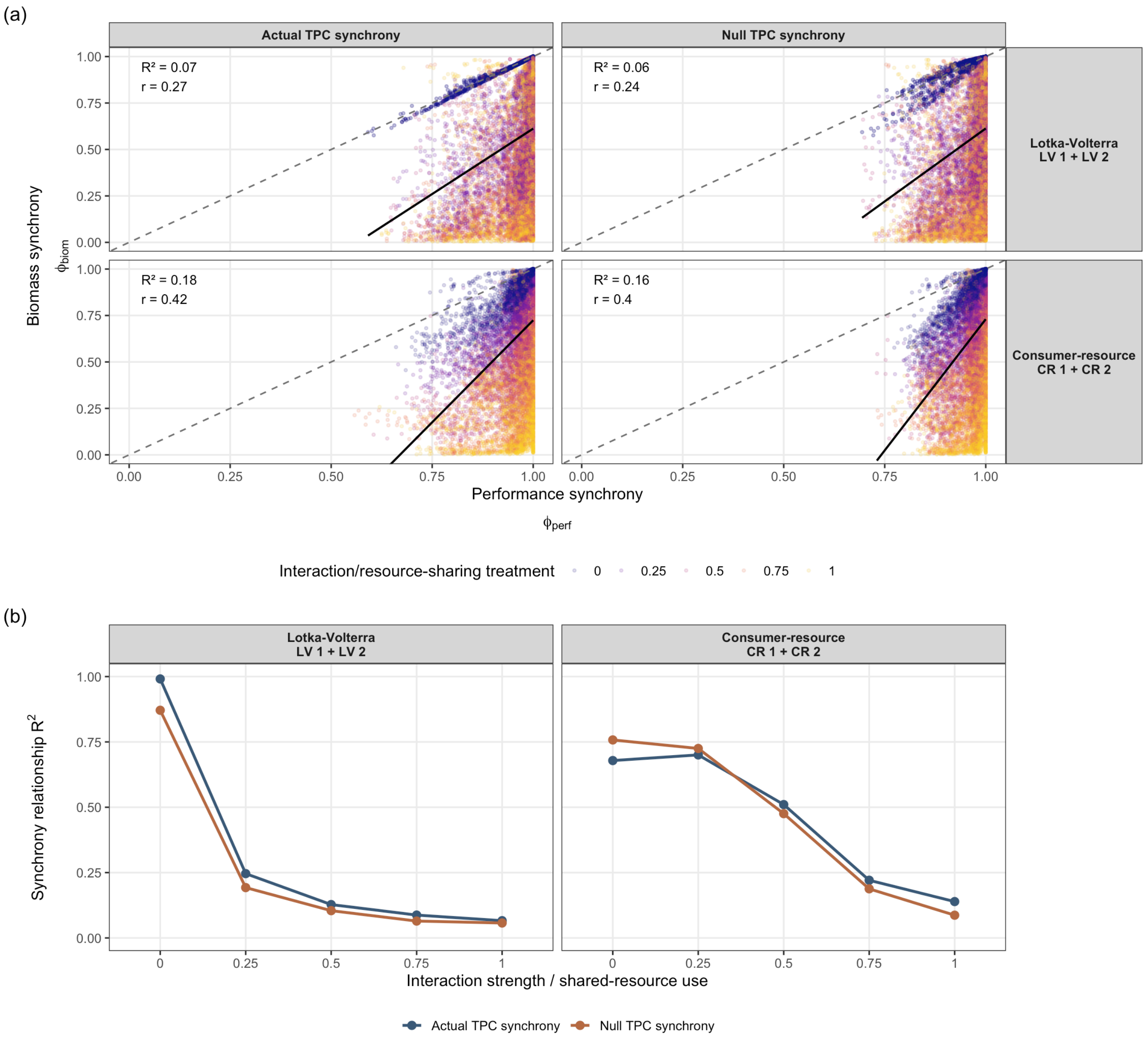


Figure S 10: Comparison between synchrony predicted from species thermal performance curves and synchrony realised in the abundance simulations. Predicted synchrony was calculated from species TPCs under either the realised temperature series or the null temperature distribution. Realised synchrony was calculated from simulated species abundances. The top panels show the overall relationship; the dashed line is the 1:1 line and the black line is a linear fit. The bottom panels show the same relationship as $R^2$ values calculated separately for each interaction-strength or shared-resource-use level.

| **framework** | **interaction/resource-sharing treatment** | **Actual TPC synchrony** | **Null TPC synchrony** |
|---|---|---|---|
| Consumer-resource | 0 | 0.679 | 0.758 |
| Consumer-resource | 0.25 | 0.700 | 0.725 |

| framework | interaction/resource-sharing treatment | Actual TPC synchrony | Null TPC synchrony |
|---|---|---|---|
| Consumer-resource | 0.5 | 0.510 | 0.476 |
| Consumer-resource | 0.75 | 0.221 | 0.188 |
| Consumer-resource | 1 | 0.139 | 0.087 |
| Lotka-Volterra | 0 | 0.991 | 0.871 |
| Lotka-Volterra | 0.25 | 0.246 | 0.193 |
| Lotka-Volterra | 0.5 | 0.128 | 0.104 |
| Lotka-Volterra | 0.75 | 0.087 | 0.065 |
| Lotka-Volterra | 1 | 0.066 | 0.057 |

Table S 4: Synchrony relationship $R^2$ by interaction strength or shared-resource use.

The synchrony comparison differed between the two modelling frameworks. In the Lotka–Volterra simulations, TPC-predicted synchrony had a weak relationship with realised abundance synchrony ((R^2 = 0.07) for actual TPC synchrony and (R^2 = 0.06) for null TPC synchrony). In the consumer-resource simulations, the relationship had higher explanatory power ((R^2 = 0.18) for actual TPC synchrony and (R^2 = 0.16) for null TPC synchrony). The consumer-resource model therefore carried more of the performance-curve synchrony signal into realised abundance synchrony. The treatment-specific (R^2) values show that this is not just an artefact of pooling across the resource-sharing gradient. Lotka–Volterra abundance synchrony was most predictable from TPC synchrony when interactions were absent, and this relationship declined as interaction strength increased. In the consumer-resource simulations, synchrony prediction had the highest explanatory power when resource use was mostly private and declined as shared-resource use increased. In both frameworks, interactions mediated how much species-level thermal synchrony was retained in realised abundance synchrony.
