## Supplementary Information 3 for "Community performance curves predict temporal stability despite interaction effects"

### 1 Scope of this supplement

This supplement provides the empirical methods, data-processing steps, and supporting analyses for the ciliate community experiment. The main manuscript reports the central empirical test of whether variability in community performance predicts community biomass variability. The material below gives the experimental design, construction of empirical thermal performance curves, calculation of null and actual CV(CP), detrending of biomass time series, model outputs, and mechanistic analyses based on performance synchrony and population variability.

### 2 Empirical study

#### 2.1 Experimental overview

The experimental study consisted of two independent experiments:

1. **Thermal performance curve experiment**: This experiment quantified species intrinsic performance along the temperature gradient (Figure S 1), yielding thermal performance curves for each species. These curves were used to calculate $CV(CP_{\text{null}})$ and $CV(CP_{\text{actual}})$ under the temperature regimes used in the community experiment (Figure S 2).

2. **Community experiment**: Selected communities were exposed to fluctuating temperature regimes. Community biomass variability was then related to $CV(CP)$ and to trait-summary predictors.

| Experiment | Duration | Sampling days | Total samples | Outcome / purpose |
|---|---|---|---|---|
| Thermal performance curve experiment | 15 days | 10 | 1,062 | Estimate species thermal performance curves and intrinsic growth rates across temperature. |
| Community experiment | 37 days | 27 | 2,916 | Test whether community performance variability predicts community biomass variability under fluctuating temperatures. |

Table S 1: Overview of the two experiments included in this report.

#### 2.2 Thermal performance curve experiment

A monoculture experiment was used to estimate intrinsic growth across temperature for the five ciliate species. The resulting thermal performance curves (TPCs) are the basis for the community performance summaries used in the community experiment.

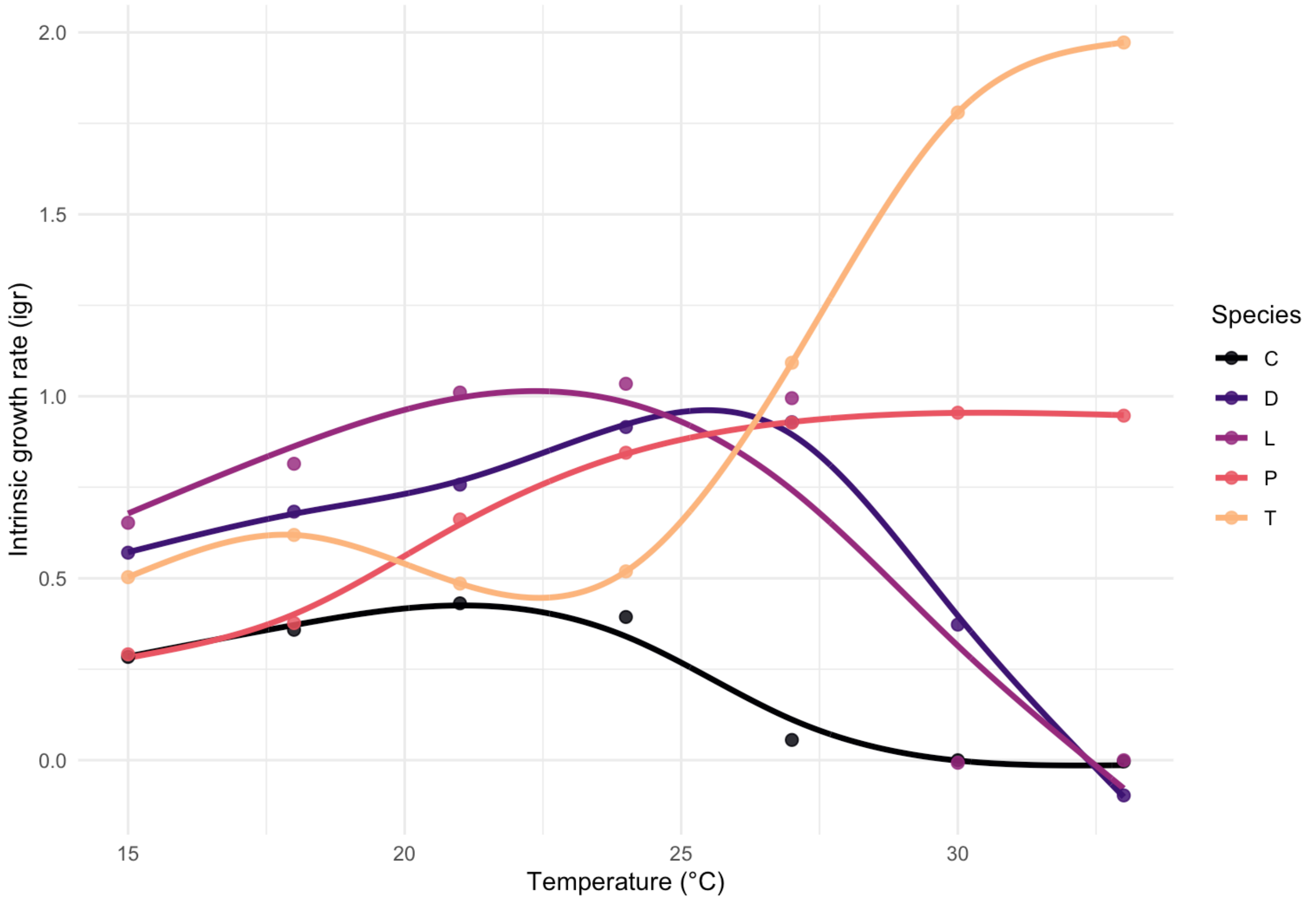


Figure S 1: Density-based thermal performance curves. Points show estimated intrinsic growth rates from the early growth window; lines show the predicted species-specific thermal performance curves used in the CP analyses.

##### 2.2.1 Temperature regime

The community experiment required predefined temperature time series to calculate $CV(CP_{\text{actual}})$. Temperature treatments consisted of four fluctuating time series. Temperature values were drawn from a normal distribution with reddened noise ($\gamma = 0.8$), a mean of 22.5°C, and a standard deviation of 3.5°C (Figure S 2). Before the fluctuating treatments began, communities were kept at a constant 22.5°C for four days; this initial stable period was later removed before analysis. The same four generated temperature time series were used as experimental temperature regimes in the community experiment and to calculate actual community performance from the species-specific TPCs.

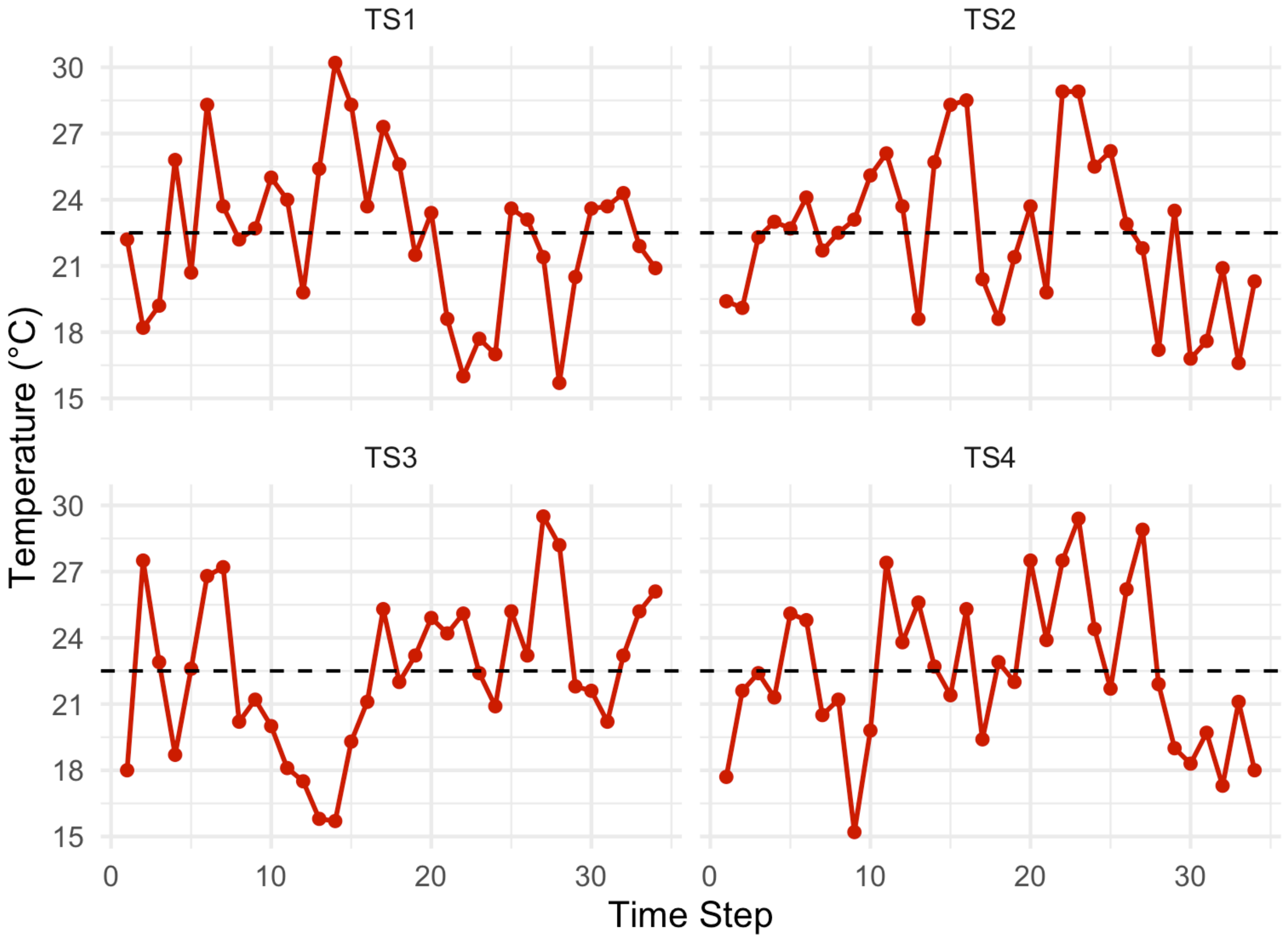


Figure S 2: Experimental temperature regimes. The four fluctuating temperature time series were generated around a mean temperature of 22.5°C, shown by the dashed horizontal line, and were later used to calculate experienced community performance.

##### 2.2.2 Actual community performance variability

The selected community compositions span low, intermediate, and high actual community performance variability within each temperature regime and richness level. Note that $CV(CP)$ was inherently related to species richness (see also Figure S 8).

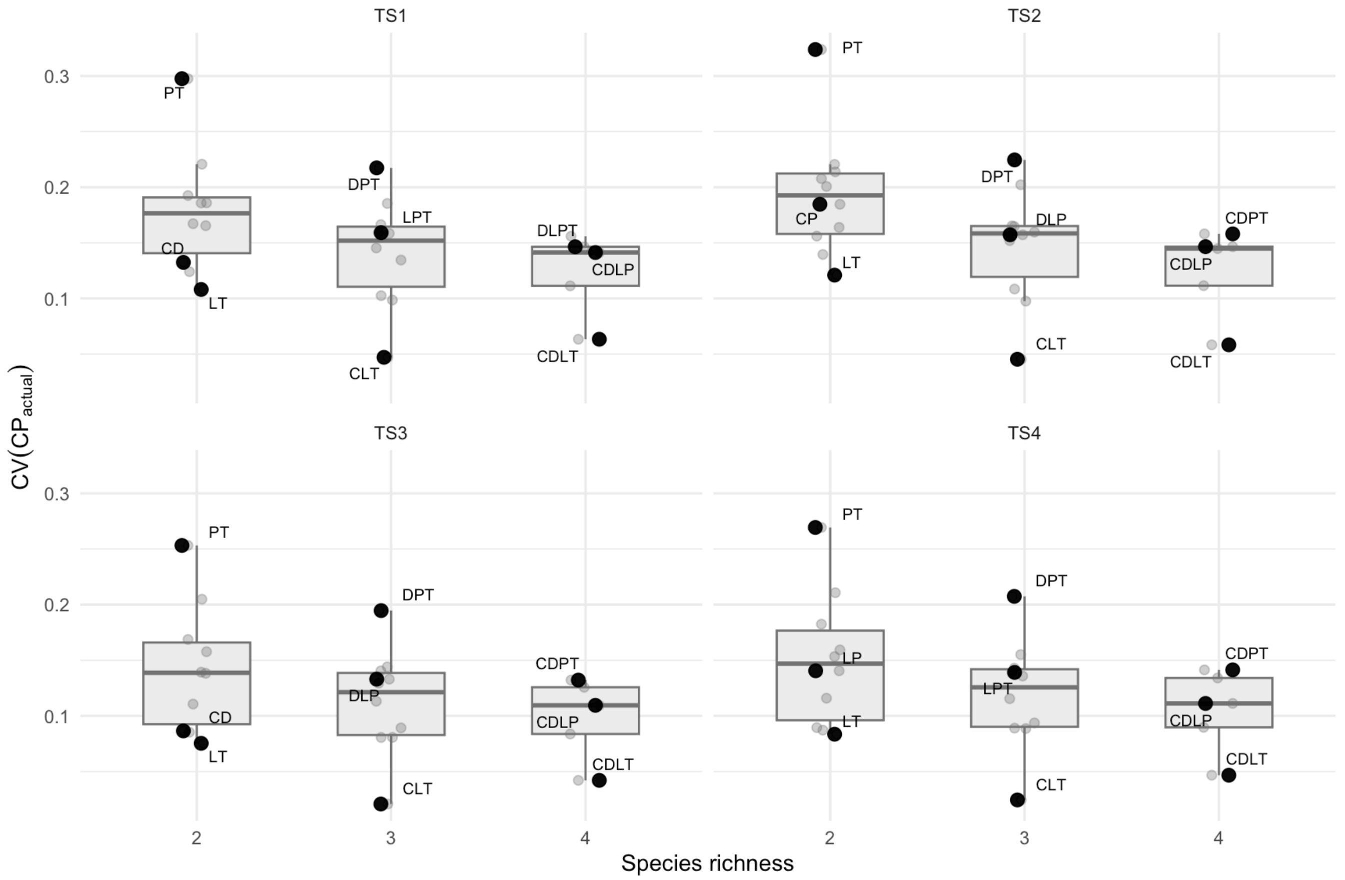


Figure S 3: Actual community performance variability under each temperature regime. Grey points show all possible species combinations not selected for the community experiment, and coloured labelled points show the selected community compositions. CV(CP) is expected to vary with species richness because CP is calculated from the summed species-level performance curves.

#### 2.3 Community experiment

Communities were assembled from the five-species ciliate pool and designed to span three richness levels: two, three, and four species. For each temperature regime and richness level, three community compositions were selected to represent a gradient in expected community performance variability: the composition with the lowest $CV(CP)$, the composition with the highest $CV(CP)$, and the composition closest to the median $CV(CP)$ (Figure S 3). This produced a factorial design with three richness levels, three selected compositions per richness level, four temperature regimes, and three replicate microcosms per treatment combination ($n = 108$). We sampled 27 times over 37 days, resulting in 2916 samples.

##### 2.3.1 Analysis

To quantify temporal variability in community biomass, the initial stable period (first two sampling days) was removed, leaving 2700 samples, of which 3 were excluded due to sampling errors. Species-level time series were then detrended before calculating community variability. For each species within each community composition and across temperature time series, a smooth generalized additive model (GAM) was fitted to describe the average change in density or biomass over experimental day (Figure S 4). This smooth trend was fitted using all replicate observations available for that species-composition combination. The fitted trend represents the shared temporal trajectory expected from gradual directional change over the experiment.

Detrended biomass was calculated by subtracting the fitted biomass trend from the observed biomass value. This leaves the short-term deviation around the smooth temporal trend, rather than the raw biomass trajectory itself. At the community level, these detrended species-level biomass values were averaged within each sample and day. Community variability was then quantified as the standard deviation of detrended community biomass across days for each sample.

Observed community variability was analysed using four predictors: variability in actual community performance ($CV(CP_{\text{actual}})$), variability in null community performance ($CV(CP_{\text{null}})$), the absolute distance between mean thermal optimum and mean environmental temperature ($|\overline{T}_{opt} - \overline{T}|$), and variation in thermal optima among species ($sd(T_{opt})$). Each predictor was analysed in a separate linear model with detrended community biomass variability as the response variable. Log-transformations were applied to improve model diagnostics. For these models, the coefficient estimate (β), *t* value, *p* value, and $R^2$ are reported.

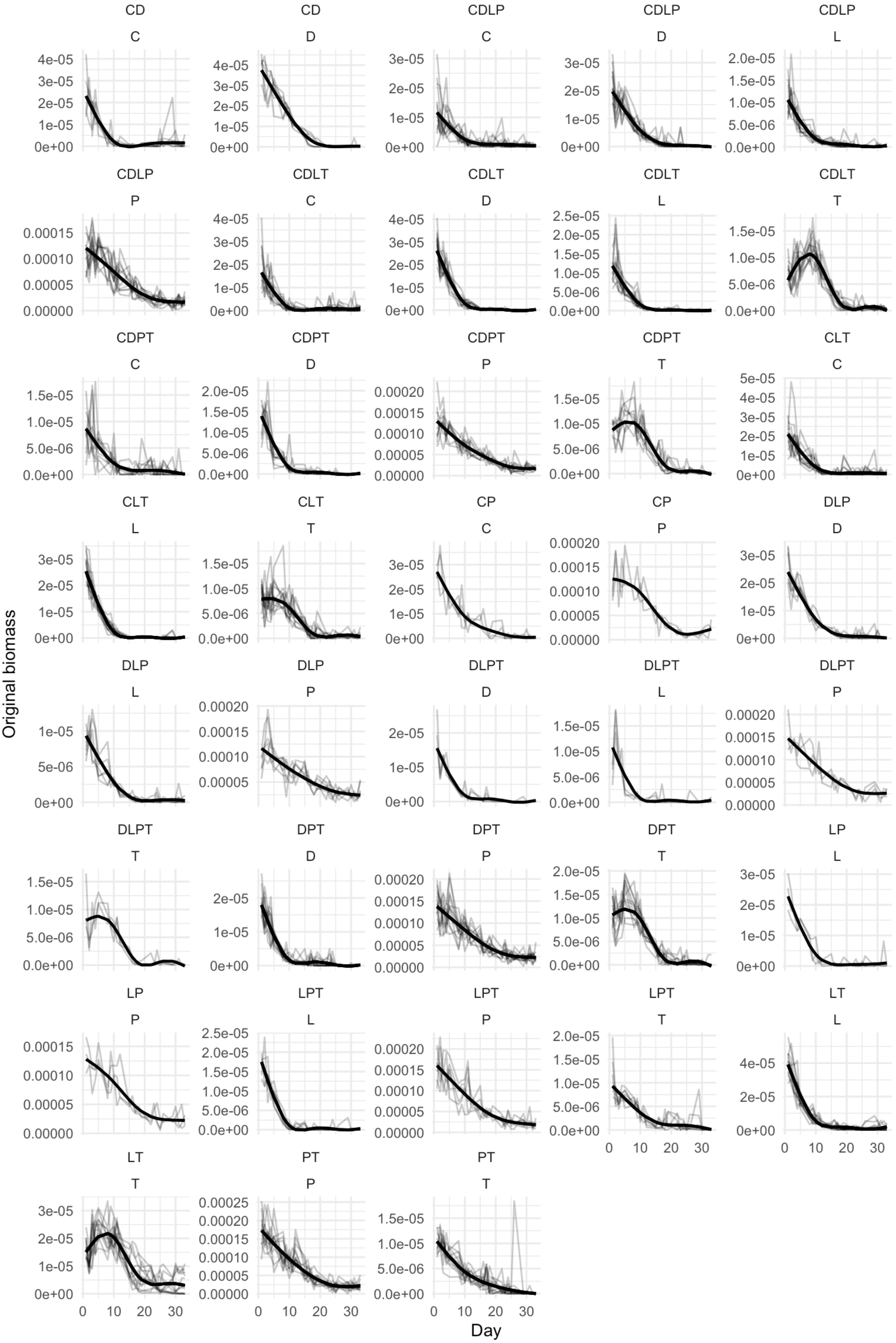
CD C
CD D
CDLP C
CDLP D
CDLP L
CDLP P
CDLT C
CDLT D
CDLT L
CDLT T
CDPT C
CDPT D
CDPT P
CDPT T
CLT C
CLT L
CLT T
CP C
CP P
DLP D
DLP L
DLP P
DLPT D
DLPT L
DLPT P
DLPT T
DPT D
DPT P
DPT T
LP L
LP P
LPT L
LPT P
LPT T
LT L
LT T
PT P
PT T
Original biomass
Day

Figure S 4: Detrending of species-level biomass time series. Thin lines show observed biomass trajectories for replicate microcosms, and thick lines show the pooled GAM trend fitted for each composition-by-species combination before calculating detrended biomass variability.

##### 2.3.2 Main Result

Variability of intrinsic community performance was positively related to variability of community biomass (Figure S 5). $CV(CP_{\text{actual}})$ explained more variation in biomass variability ($R^2 =$ 0.67) than $CV(CP_{\text{null}})$ ($R^2 =$ 0.40). Communities with higher $CV(CP_{\text{actual/null}})$ had higher temporal biomass variability ($\beta_{actual}$ = 9.25e-01, *p* = < 0.001; $\beta_{null}$ = 8.59e-01, *p* = < 0.001).

Similarly, trait summaries were also predictive of stability, with $|\overline{T}_{opt} - \overline{T}|$ explaining less variance ( $R^2 =$ 0.25) than $sd(T_{opt})$ ($R^2 =$ 0.27). The larger the distance between the average temperature and the average thermal optimum was (large $|\overline{T}_{opt} - \overline{T}|$), the more variable the communities were (β = 6.98e-02, *p* = < 0.001). On the other hand, the spread of the thermal optima ($sd(T_{opt})$) had a stabilising effect ( β = -1.03e-01, *p* = < 0.001).

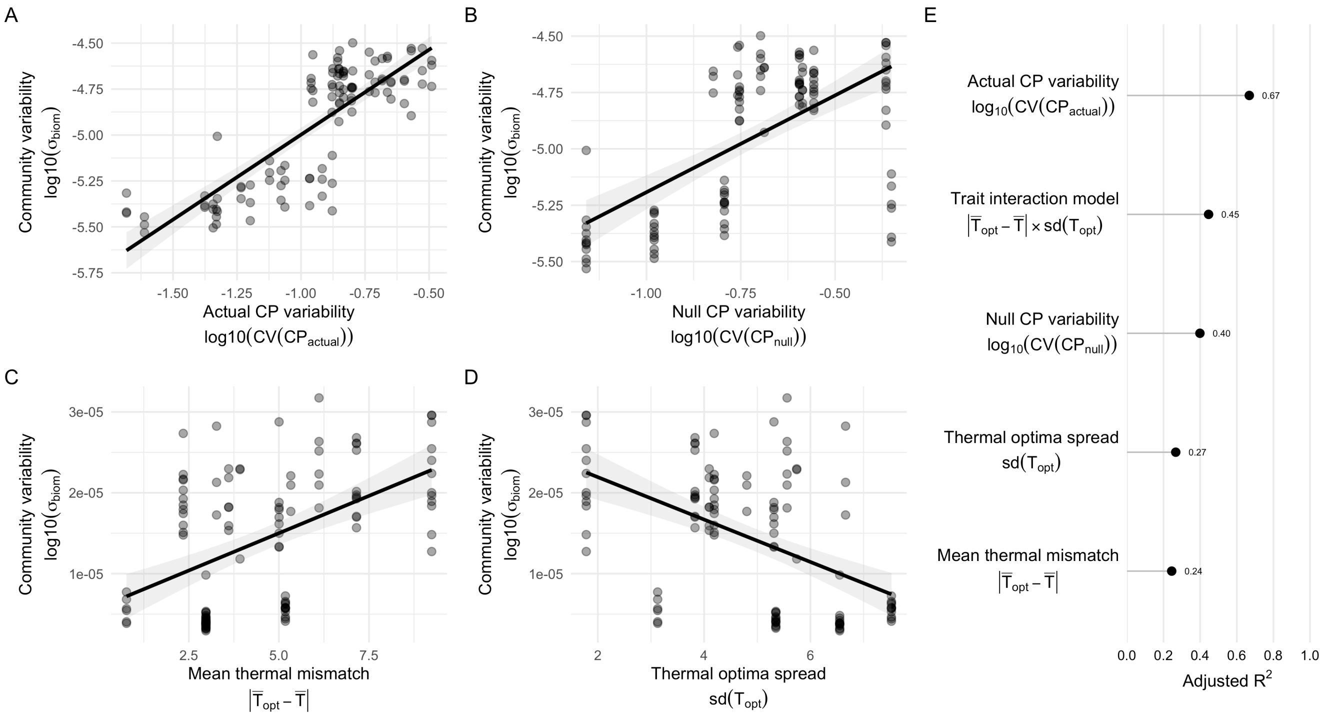


Figure S 5: Predictors of detrended community biomass variability. Panels A–D show the relationships between community biomass variability and (A) actual community performance (CP) variability, (B) null CP variability, (C) the absolute difference between mean species thermal optimum and mean experienced temperature, and (D) variation in species thermal optima. Solid lines show fitted linear regressions and shaded areas show 95% confidence intervals. Panel E compares the R^2values of the four single-predictor models with that of the trait-summary model, which includes mean thermal mismatch, thermal-optimum variation, and their interaction.

##### 2.3.3 Full Model Results

The full model included $CV(CP_{\text{actual}})$, species richness, temperature time series, and the interaction between $CV(CP_{\text{actual}})$ and richness. Together these terms explained most of the variation in detrended community biomass variability (adjusted $R^2$ = 0.83; overall $F$(8, 99) = 67.11, *p* = < 0.001). $CV(CP_{\text{actual}})$ accounted for a substantial component of model fit ($F$(1, 99) = 426.97, *p* = < 0.001), and the relationship between $CV(CP_{\text{actual}})$ and biomass variability differed among richness levels ($F$(2, 99) = 12.03, $p$ = < 0.001).

Estimated slopes were positive at all richness levels (richness 2: β = 1.26; richness 3: β = 0.92; richness 4: β = 1.49). The slope was shallower for three-species communities than for two-species communities, while the four-species slope was steeper but had overlapping uncertainty with the two-species slope (richness 3 vs. richness 2: β = -3.42e-01, $p$ = 0.005; richness 4 vs. richness 2: β = 2.31e-01, $p$ = 0.126). Temperature time series also contributed to the full model ($F$(3, 99) = 6.42, $p$ = < 0.001). Figure S 6 shows the $CV(CP_{\text{actual}})$ relationship grouped by richness and by temperature time series.

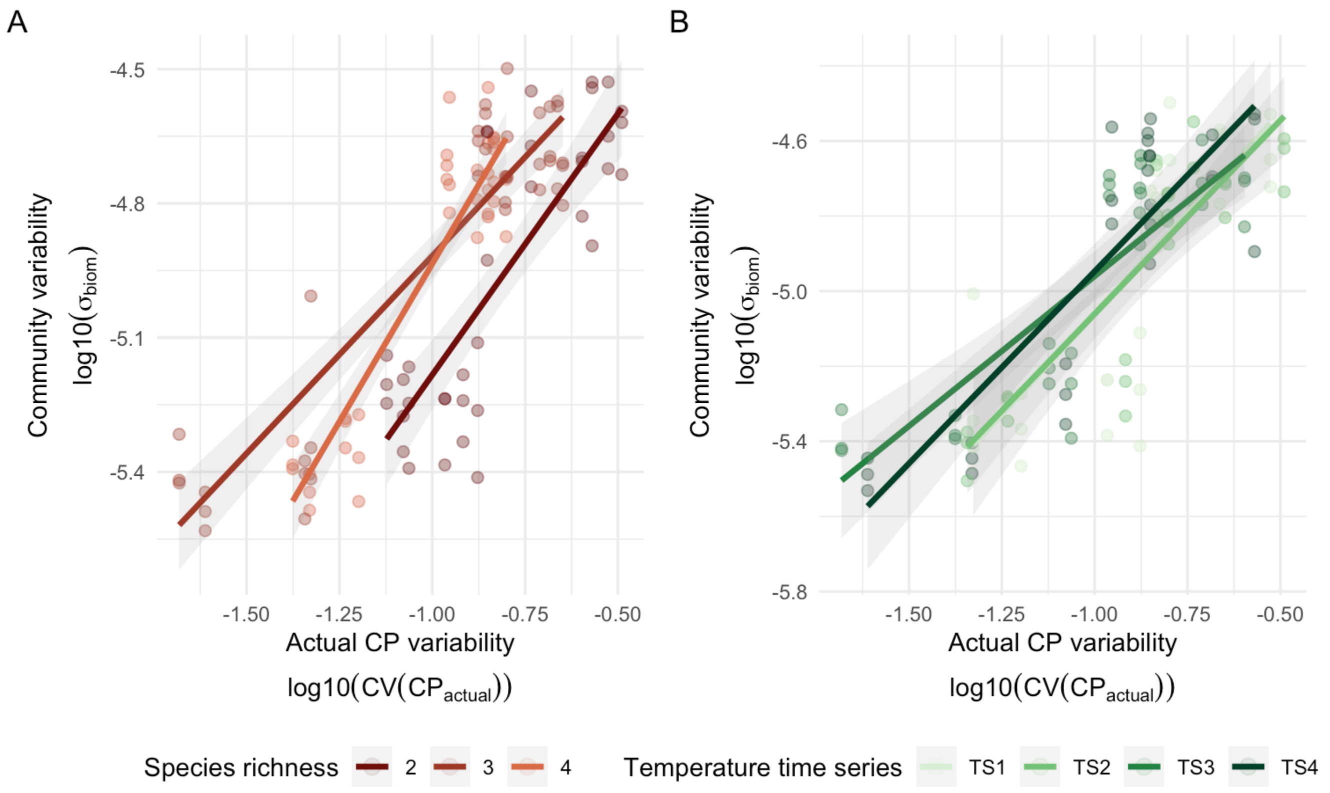


Figure S 6: Full-model view of the relationship between actual community performance variability and detrended community biomass variability. The same relationship is shown by species richness on the left and by temperature time series on the right, with points showing communities and lines showing fitted linear trends.

#### 2.3.4 Mechanistic breakdown

For species biomass time series $b_i(t)$, community biomass is $B(t) = \sum_i b_i(t)$. The coefficient of variation of community biomass can be decomposed into biomass synchrony ($\phi_{\text{biom}}$) and weighted mean population variability:

$$CV_{\text{biom}} = \frac{\sigma(B(t))}{\bar{B}(t)} = \sqrt{\phi_{\text{biom}}} \sum_i w_i CV(b_i(t)), \qquad (1)$$

where

$$\phi_{\text{biom}} = \frac{\sigma^2\left(\sum_i b_i(t)\right)}{\left(\sum_i \sigma(b_i(t))\right)^2} \quad \text{and} \quad w_i = \frac{\bar{b_i}(t)}{\bar{B}(t)}. \qquad (2)$$

Because the biomass time series were detrended and the response is the standard deviation of detrended community biomass rather than raw biomass $CV$, the empirical relationship differs from Equation 1. The response is $\sigma_{\text{biom}}$, which can be written as

$$\sigma_{\text{biom}} = \sqrt{\phi_{\text{biom}}} \sum_i \sigma(b_i(t)) = \sqrt{\phi_{\text{biom}}}\, S \bar{\sigma}_{\text{biom}}, \qquad (3)$$

where mean population biomass variability is defined as

$$\bar{\sigma}_{\text{biom}} = \frac{1}{S} \sum_i \sigma(b_i), \qquad (4)$$

and $S$ is species richness. Analogously, for species performance $r_i(t)$, community performance is $R(t) = \sum_i r_i(t)$ and

$$CV(CP_{\text{actual}}) = \frac{\sigma(R(t))}{\bar{R}(t)} = \sqrt{\phi_{\text{perf}}}\, \tilde{CV}_{perf}, \qquad (5)$$

where performance synchrony is defined as

$$\phi_{\text{perf}} = \sigma^2(R(t))/(\sum_i \sigma(r_i(t)))^2, \qquad (6)$$

and mean species performance variability as

$$\tilde{CV}_{perf} = \sum_i w_i CV(r_i(t)), \qquad (7)$$

with $w_i = \bar{r}_i(t)/\bar{R}(t)$. Thus, $CV(CP_{\text{actual}})$ integrates both performance synchrony ($\phi_{\text{perf}}$) and mean species performance variability ($\tilde{CV}_{perf}$). This motivates tests of whether performance synchrony predicts biomass synchrony and whether mean species performance variability predicts mean population biomass variability (Figure S 7):

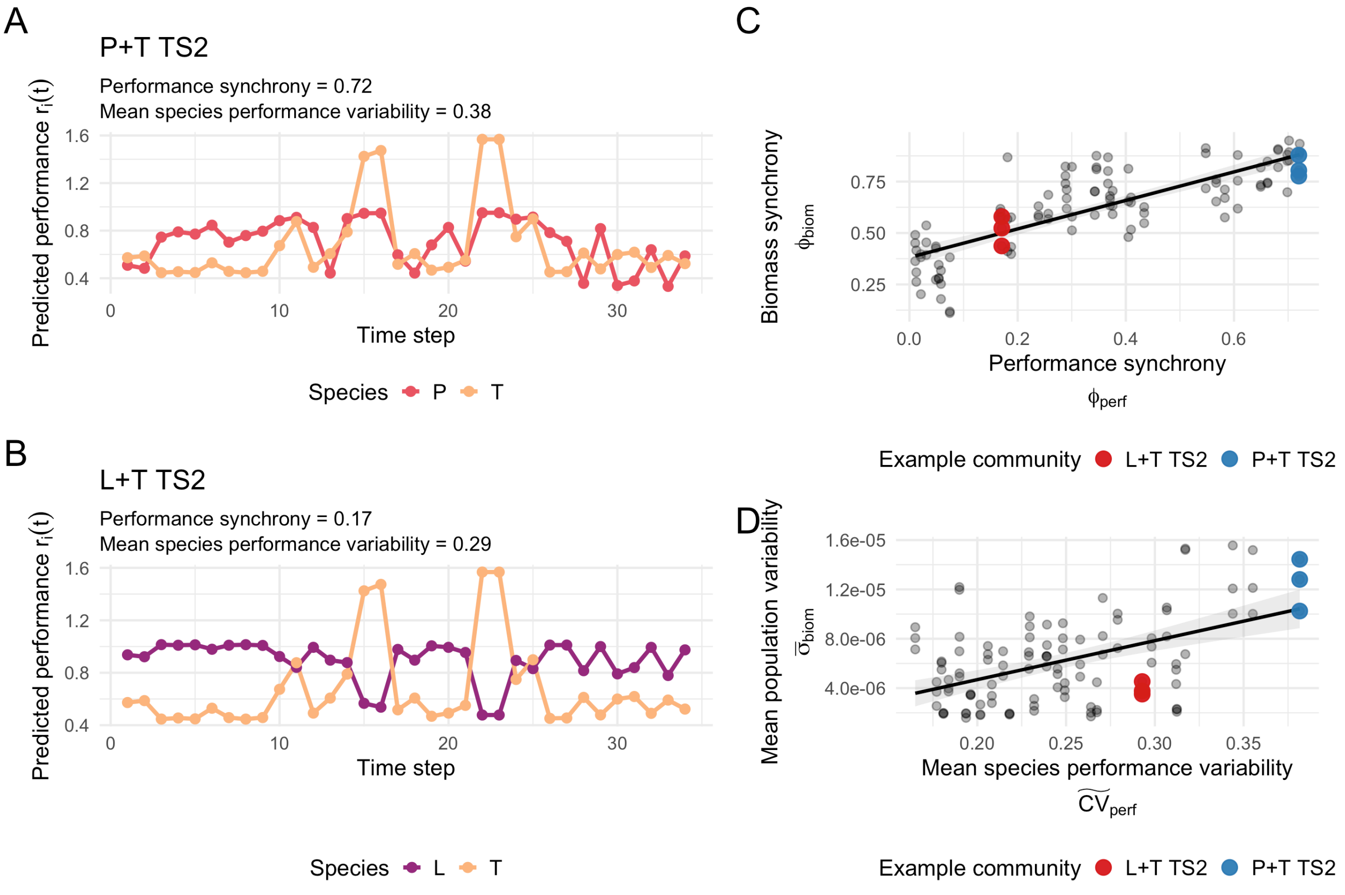


Figure S 7: Mechanistic breakdown of performance variability. The left panels show predicted species performance $r_i(t)$ through TS2 for two example communities, P+T and L+T, using the species thermal performance GAMs. The right panels show how performance synchrony and mean species performance variability relate to biomass synchrony and mean population variability across all communities; larger coloured points mark the two example communities.

Across all communities, performance synchrony was positively related to synchrony of biomass fluctuations (Figure S 7, panel C): communities with more synchronous performance also showed more synchronous biomass dynamics ($R^2 =$ 0.65; β = 6.93e-01, *p* = < 0.001). Mean species performance variability was positively related to mean population variability (Figure S 7, panel D) ( $R^2 =$ 0.24; β = 3.16e-05, *p* = < 0.001).

Structural equation modelling (SEM) was used to test whether $CV(CP_{\text{actual}})$ was related to community variability through synchrony and mean population variability, while accounting for species richness. The model also included the association between species richness and intrinsic performance variability ($CV(CP_{\text{actual}})$). All variables were standardized; biomass synchrony, biomass variability terms, and richness were log-transformed so that the interactive effect between synchrony and mean population variability changed to additive.

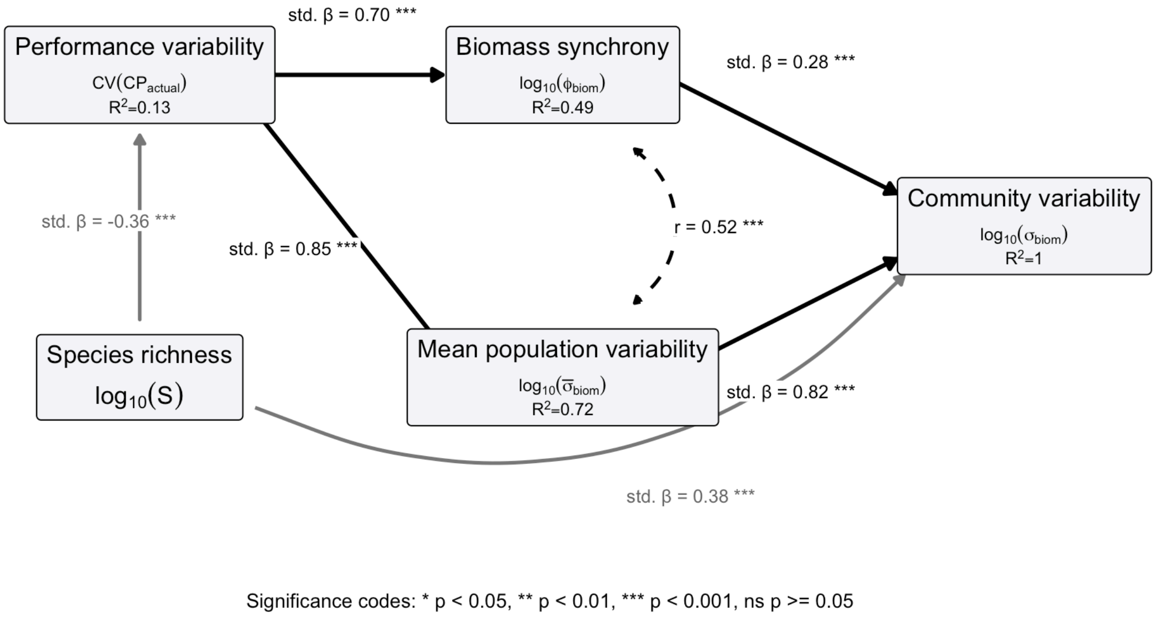


Figure S 8: Structural equation model for the mechanistic pathways from performance variability to community biomass variability. Path labels show standardized effects and significance codes, and node labels show the variable name, mathematical expression, and explained variance. Grey arrows show richness paths, and the dashed double-headed arrow shows the residual correlation between biomass synchrony and mean population variability.

In the SEM, $CV(CP_{\text{actual}})$ was associated with both $\log_{10}$ biomass synchrony (β = 0.70, z = 9.39, *p* = < 0.001; $R^2$ = 0.49) and $\log_{10}$ mean population biomass variability (β = 0.85, z = 16.55, *p* = < 0.001; $R^2$ = 0.72) (Figure S 8). Downstream, both biomass components were related to $\log_{10}(\sigma_{\text{biom}})$, with a larger standardized coefficient for the mean-population-variability pathway (β = 0.82, z = 308.90, *p* = < 0.001) than for the synchrony pathway (β = 0.28, z = 112.53, *p* = < 0.001). Richness was related to $CV(CP_{\text{actual}})$ (β = -0.36, z = -3.93, *p* = < 0.001; $R^2$ = 0.13). Together, the model explained all variation of $\log_{10}(\sigma_{\text{biom}})$ ($R^2$ = 1.00) due to the mathematically exact relationship (see Equation 3). Model fit statistics were $\chi^2$(3) = 5.71, *p* = 0.127, CFI = 0.998, RMSEA = 0.091, and SRMR = 0.031.
