## Supplementary Information 4 for "Community performance curves predict temporal stability despite interaction effects"

PUBLISHED
October 6, 2026

#### 1 Overview

This derivation provides a mechanistic reason that temporal variation in CP should contain information about temporal variation in total community biomass, together with the conditions that make this prediction closer to a calibrated equality.

The derivation shows that:

$$\mathrm{CV}_B^2 \approx \frac{\mathrm{CV}_{CP}^2 - (1-p)\mathrm{CV}_{\mathrm{sp}}^2}{p}. \qquad (1)$$

In this equation:

- $\mathrm{CV}_B^2$ is the squared coefficient of variation of total community abundance. This is the square of the **community-level variability**, a common measure of **the inverse of community stability**.
- $\mathrm{CV}_{CP}^2$ is the squared coefficient of variation of community performance. This is the square of **the response-diversity measure** (CV CP) calculated from the species performance curves.
- $\mathrm{CV}_{\mathrm{sp}}^2$ is the weighted squared coefficient of variation of species abundances:

$$\mathrm{CV}_{\mathrm{sp}}^2 = \sum_{i=1}^{m} \pi_i \mathrm{CV}^2(N_i)$$

and

$$\pi_i = \frac{\langle N_i \rangle^2}{\sum_{j=1}^{m} \langle N_j \rangle^2}$$

where $m$ is the number of species in the community, and $N_i$ is the abundance of species $i$.

- $p$ is a weight that depends on the mean structure and heterogeneity of the interaction matrix. When interaction heterogeneity is small relative to the average interaction structure, $p$ is close to one. In contrast, when interaction heterogeneity is large relative to the average interaction structure, $p$ is smaller.

Therefore, interspecific interactions, via $p$, control how strongly species-level abundance variability contributes to the relationship between CP variability and total community abundance variability. When pair-specific interaction heterogeneity is small, $p$ is close to one and the relationship between $\mathrm{CV}_{CP}^2$ and $\mathrm{CV}_B^2$ is nearly one-to-one. When pair-specific interaction heterogeneity is large, $p$ is smaller and species-level abundance variability contributes more strongly.

Thus, if an interaction matrix is well approximated by its mean structure, the variability of community performance should be a good predictor of the variability of total community abundance. If the interaction matrix is highly heterogeneous, species-level abundance variability should play a larger role.

The derivation follows. It is conditional on positive and bounded population trajectories, weak immigration, separation between environmental and population-dynamic timescales, and a statistical approximation for the interaction matrix.

The derivation is followed by connection to the LV publication experiments, in particular showing the interaction distributions in the experiments, and the realised values of $p$ in the simulated communities. We show that in the simulations even when interactions and their effects on population dynamics are strong, the realised $p$ values are high, meaning that the average interaction structure dominates over pair-specific heterogeneity. This is consistent with the strong CP–biomass relationship observed in the simulations.

### 2 Temporal variability of an aggregate quantity

This section establishes useful terms for describing variability. Ecologically, community performance and total biomass are both aggregate quantities: they combine contributions from several species. Before asking why CP should predict biomass stability, we need a way to express the variability of a community-level sum in terms of the variability and covariation of its species-level parts.

For a time-dependent quantity $X(t)$ observed over an interval of length $\mathcal{T}$, define the temporal expectation

$$\langle X^q \rangle = \frac{1}{\mathcal{T}} \int_0^{\mathcal{T}} X(t)^q \, \mathrm{d}t. \qquad (2)$$

The squared coefficient of variation is

$$\mathrm{CV}^2(X) = \frac{\mathrm{Var}(X)}{\langle X \rangle^2} = \frac{\langle X^2 \rangle}{\langle X \rangle^2} - 1. \qquad (3)$$

Suppose that $X(t)$ is the sum of $m$ species-level quantities $y_i(t)$:

$$X(t) = \sum_{i=1}^{m} y_i(t). \qquad (4)$$

Its first two temporal moments are

$$\langle X \rangle = \sum_{i=1}^{m} \langle y_i \rangle \qquad (5)$$

and

$$\langle X^2 \rangle = \sum_{i=1}^{m} \sum_{j=1}^{m} \langle y_i y_j \rangle. \qquad (6)$$

Let $\mathbf{y} = (y_1, \ldots, y_m)^\top$ and let $\mathbf{1}$ denote an $m$-vector of ones. It follows that

$$\mathrm{CV}^2(X) = \frac{\mathbf{1}^\top \langle \mathbf{y}\mathbf{y}^\top \rangle \mathbf{1}}{\mathbf{1}^\top \langle \mathbf{y} \rangle \langle \mathbf{y} \rangle^\top \mathbf{1}} - 1. \qquad (7)$$

The same expression is obtained if $X$ is defined as a species mean because multiplying a time series by the constant $1/m$ does not change its coefficient of variation.

Applying Equation 7 with $y_i(t) = \tilde{r}_i(T_t)$ gives

$$\mathrm{CV}_{CP}^2 = \frac{\mathbf{1}^\top \langle \tilde{\mathbf{r}}\tilde{\mathbf{r}}^\top \rangle \mathbf{1}}{\mathbf{1}^\top \langle \tilde{\mathbf{r}} \rangle \langle \tilde{\mathbf{r}} \rangle^\top \mathbf{1}} - 1. \qquad (8)$$

Thus, CP variability depends on both the temporal variances of individual species' performances and their temporal covariances.

### 3 Relationship to the continuous-time LV simulation model

This section connects the statistical quantity CP to the population dynamics used in the simulations. The aim is to show that, in the continuous-time LV model, CP is not an external summary imposed after the fact: it is built from the same temperature-dependent growth-rate terms that drive changes in abundance.

For species $i$, the simulation model is

$$\frac{\mathrm{d}N_i}{\mathrm{d}t} = N_i(t)\tilde{r}_i(T_t)\left[1 - \frac{\sum_{j=1}^{m} \alpha_{ij} N_j(t)}{K_i(T_t)}\right] + I, \qquad (9)$$

where

$$\tilde{r}_i(T_t) = b_i(T_t) - d_i(T_t) + \epsilon \qquad (10)$$

and

$$K_i(T_t) = \frac{\tilde{r}_i(T_t)}{\beta + \delta}. \qquad (11)$$

Substitution of Equation 11 into Equation 9 gives

$$\frac{\mathrm{d}N_i}{\mathrm{d}t} = N_i(t)\left[\tilde{r}_i(T_t) - (\beta + \delta)\sum_{j=1}^{m} \alpha_{ij} N_j(t)\right] + I. \qquad (12)$$

Defining the effective interaction coefficients

$$A_{ij} = (\beta + \delta)\alpha_{ij} \qquad (13)$$

produces

$$\frac{\mathrm{d}N_i}{\mathrm{d}t} = N_i(t)\left[\tilde{r}_i(T_t) - \sum_{j=1}^{m} A_{ij} N_j(t)\right] + I. \qquad (14)$$

Thus, apart from immigration, CP is constructed from the environmental growth-rate terms that directly drive abundance dynamics. Under Equation 11, the temperature dependence assigned to $K_i$ does not create a second independent pathway: it yields the fixed effective interaction coefficients in Equation 13.

### 4 Long-run mean relationship

This section explains why persistent populations must, on average, balance growth against interaction pressure. This is useful because it turns the dynamic LV equation into a relationship between mean environmental performance and mean abundances, giving the first bridge from species performance curves to community biomass.

We first consider $I = 0$. For positive trajectories, division of Equation 14 by $N_i$ gives

$$\frac{\mathrm{d}\log N_i}{\mathrm{d}t} = \tilde{r}_i(T_t) - \sum_{j=1}^{m} A_{ij} N_j(t). \qquad (15)$$

Averaging over $[0, \mathcal{T}]$ yields

$$\langle \tilde{r}_i \rangle - \sum_{j=1}^{m} A_{ij} \langle N_j \rangle = \frac{\log N_i(\mathcal{T}) - \log N_i(0)}{\mathcal{T}}. \qquad (16)$$

If abundances remain bounded above and bounded away from zero over the averaging interval, then there exist constants $(N_{\min} > 0)$ and $(N_{\max} < \infty)$ such that

$$N_{\min} \leq N_i(t) \leq N_{\max}.$$

The right-hand side of Equation 16 is a boundary term: it depends only on the abundance at the start and end of the averaging interval. If abundances remain bounded in this way, then the total endpoint change in log abundance is also bounded:

$$\left| \log N_i(\mathcal{T}) - \log N_i(0) \right| \leq \log\left(\frac{N_{\max}}{N_{\min}}\right).$$

A bounded endpoint change divided by an increasingly long averaging interval must become negligible. It therefore follows that

$$\frac{\log N_i(\mathcal{T}) - \log N_i(0)}{\mathcal{T}} \to 0 \qquad \text{as} \qquad \mathcal{T} \to \infty.$$

Thus, over a long interval, the average log-growth rate is approximately zero for a bounded persistent population. This is the key step in the derivation: persistence converts the dynamic LV equation into an approximate balance between the mean intrinsic growth rates and the mean competitive effects:

$$\left\langle \tilde{r}_i \right\rangle \approx \sum_{j=1}^{m} A_{ij} \left\langle N_j \right\rangle.$$

or equivalently,

$$\langle \tilde{\mathbf{r}} \rangle \approx \mathbf{A} \langle \mathbf{N} \rangle. \qquad (17)$$

With continuous immigration, the corresponding identity is

$$\langle \tilde{r}_i \rangle - \sum_{j=1}^{m} A_{ij} \langle N_j \rangle + I \left\langle \frac{1}{N_i} \right\rangle = \frac{\log N_i(\mathcal{T}) - \log N_i(0)}{\mathcal{T}}. \qquad (18)$$

Therefore, Equation 17 is accurate when immigration is absent or when $I\langle 1/N_i \rangle$ is small relative to the other terms. Immigration may be especially consequential for species maintained at very low abundance.

### 5 Slow-environment approximation

This section adds the ecological assumption that temperature changes slowly enough for the community to remain near its moving equilibrium. Under that assumption, abundance variation can be approximated from the temperature-driven growth-rate vector, allowing CP variability and biomass variability to be compared through the interaction matrix.

Assume that temperature changes slowly relative to the intrinsic relaxation of the biological dynamics. The abundance vector then remains close to a temperature-dependent quasi-equilibrium $\mathbf{N}^*(T)$ satisfying, when immigration is negligible,

$$\tilde{\mathbf{r}}(T) \approx \mathbf{A}\mathbf{N}^*(T). \qquad (19)$$

Equivalently,

$$\mathbf{N}^*(T) \approx \mathbf{A}^{-1}\tilde{\mathbf{r}}(T), \qquad (20)$$

when $\mathbf{A}$ is invertible and the equilibrium is feasible. Consequently,

$$\langle \tilde{\mathbf{r}}\tilde{\mathbf{r}}^\top \rangle \approx \mathbf{A}\langle \mathbf{N}^*\mathbf{N}^{*\top}\rangle \mathbf{A}^\top. \qquad (21)$$

For notational simplicity, we henceforth write $\mathbf{N}$ for the quasi-equilibrium vector. Substitution into Equation 8 gives

$$\mathrm{CV}_{CP}^2 \approx \frac{\mathbf{1}^\top \mathbf{A}\langle \mathbf{N}\mathbf{N}^\top\rangle \mathbf{A}^\top \mathbf{1}}{\mathbf{1}^\top \mathbf{A}\langle \mathbf{N}\rangle\langle \mathbf{N}\rangle^\top \mathbf{A}^\top \mathbf{1}} - 1. \qquad (22)$$

This relationship shows that CP variability is linked to abundance variability, but that the mapping depends on the interaction matrix.

### 6 Average interaction structure and pair-specific deviations

This section asks how the interaction matrix changes the CP-biomass relationship. The ecological idea is that interactions have both an average component, shared across many species pairs, and idiosyncratic pair-specific deviations. Separating these parts lets us see when interactions mostly transmit the aggregate CP signal and when interaction heterogeneity adds species-level noise.

To obtain a more interpretable expression, assume that effective self-interactions are fixed at $a_0$ and that off-diagonal interactions have mean $\bar{a}$ and variance $\sigma_a^2$. Write

$$\mathbf{A} = (a_0 - \bar{a})\mathbf{I} + \bar{a}\mathbf{1}\mathbf{1}^\top + \mathbf{\Delta}. \qquad (23)$$

This decomposition separates the interaction matrix into two conceptually distinct parts:

1. **Average interaction structure.** The matrix $(a_0 - \bar{a})\mathbf{I} + \bar{a}\mathbf{1}\mathbf{1}^\top$ is the interaction matrix that would be obtained if every off-diagonal interaction equalled the mean interaction strength $\bar{a}$. It retains the fixed self-interaction $a_0$ and the mean effect of the other $m - 1$ species.
2. **Pair-specific deviations.** The matrix $\mathbf{\Delta}$ contains the deviations of individual interactions from that average matrix. Its off-diagonal entries have mean zero and variance $\sigma_a^2$, while its diagonal entries are zero. Thus, $\sigma_a^2$ measures interaction heterogeneity around the average interaction structure.

Define the average contribution to a row or column sum as

$$c = a_0 + (m-1)\bar{a}. \qquad (24)$$

The quantity $c$ summarizes the fixed self-interaction together with the mean contribution of the other $m-1$ species. It follows that

$$\mathbf{1}^\top \mathbf{A} = c\mathbf{1}^\top + \mathbf{1}^\top \mathbf{\Delta}. \qquad (25)$$

For an arbitrary matrix $\mathbf{Q}$, neglecting mean-zero cross terms and replacing the quadratic contribution of the deviations by its ensemble expectation gives

$$\mathbf{1}^\top \mathbf{A}\mathbf{Q}\mathbf{A}^\top \mathbf{1} \approx c^2 \mathbf{1}^\top \mathbf{Q}\mathbf{1} + m\sigma_a^2 \mathrm{tr}(\mathbf{Q}). \qquad (26)$$

The first term is the contribution expected from the average interaction structure. The second term is the expected contribution of pair-specific heterogeneity. The approximation is an ensemble statement: its accuracy for one finite interaction matrix depends on species richness, the distribution and dependence of interaction coefficients, and the extent to which the omitted terms cancel.

Applying Equation 26 to Equation 22 gives

$$\mathrm{CV}_{CP}^2 \approx \frac{c^2 \left\langle \left( \sum_i N_i \right)^2 \right\rangle + m\sigma_a^2 \sum_i \langle N_i^2 \rangle}{c^2 \left( \sum_i \langle N_i \rangle \right)^2 + m\sigma_a^2 \sum_i \langle N_i \rangle^2} - 1. \qquad (27)$$

### 7 Weighted decomposition of variability

This section collects the preceding pieces into the main prediction. It rewrites CP variability as a weighted combination of total community biomass variability and weighted species-level abundance variability, with the weight $p$ measuring how strongly the average interaction structure dominates over pair-specific heterogeneity.

Define total community abundance as

$$B(t) = \sum_{i=1}^{m} N_i(t) \qquad (28)$$

and

$$\mathrm{CV}_B^2 = \frac{\langle B^2 \rangle}{\langle B \rangle^2} - 1. \qquad (29)$$

Define weighted species-level variability as

$$\mathrm{CV}_{\mathrm{sp}}^2 = \frac{\sum_i \langle N_i^2 \rangle}{\sum_i \langle N_i \rangle^2} - 1 = \frac{\sum_i \langle N_i \rangle^2 \mathrm{CV}^2(N_i)}{\sum_i \langle N_i \rangle^2}. \qquad (30)$$

Now define

$$p = \frac{c^2 \left( \sum_i \langle N_i \rangle \right)^2}{c^2 \left( \sum_i \langle N_i \rangle \right)^2 + m\sigma_a^2 \sum_i \langle N_i \rangle^2}. \qquad (31)$$

Equation 27 becomes

$$\mathrm{CV}_{CP}^2 \approx p\mathrm{CV}_B^2 + (1-p)\mathrm{CV}_{\mathrm{sp}}^2. \qquad (32)$$

The weight $p$ compares the contribution of the average interaction structure with the contribution of pair-specific interaction heterogeneity. A large value of $p$ means that the average interaction term dominates the approximation; a smaller value means that variation among individual pairwise interactions makes a larger contribution.

For $p > 0$, the relationship can be rearranged as

$$\mathrm{CV}_B^2 \approx \frac{\mathrm{CV}_{CP}^2 - (1-p)\mathrm{CV}_{\mathrm{sp}}^2}{p}. \qquad (33)$$

Thus, a near one-to-one relationship between CP variability and total-abundance variability is expected when $p$ is close to one.

### 8 Equal-mean special case

This section gives a simpler interpretation of $p$ for the special case where species have similar mean abundances. The purpose is not to assume that communities always have equal abundances, but to make the ecological controls on $p$ easier to see: mean interaction structure pushes $p$ upward, while heterogeneity among pairwise interactions pushes it downward.

If all species have approximately the same mean abundance, $\langle N_i \rangle \approx \mu$, then

$$\left(\sum_i \langle N_i \rangle\right)^2 \approx m^2\mu^2, \qquad \sum_i \langle N_i \rangle^2 \approx m\mu^2,$$

and

$$p \approx \frac{c^2}{c^2 + \sigma_a^2}. \qquad (34)$$

When effective self-interactions are standardized to one, $a_0 = 1$, this becomes

$$p \approx \frac{[1 + (m-1)\overline{a}]^2}{[1 + (m-1)\overline{a}]^2 + \sigma_a^2}. \qquad (35)$$

Therefore, $p$ approaches one when the squared average interaction term, $[a_0 + (m-1)\overline{a}]^2$, is large relative to the variance in pairwise interaction strengths, $\sigma_a^2$. In ecological terms, the interaction matrix is then well summarized by its mean structure, and pair-specific deviations contribute relatively little. Conversely, large interaction heterogeneity reduces $p$ and increases the contribution of weighted species-level variability.

Non-negative mean abundances do not by themselves imply $p > 1/2$. The relevant condition is

$$c^2\left(\sum_i \langle N_i \rangle\right)^2 > m\sigma_a^2 \sum_i \langle N_i \rangle^2. \qquad (36)$$

### 9 Connection to the simulation results

The LV publication experiments use two different ways of constructing competitive interaction matrices. LV1 uses asymmetric uniform competition: interspecific interaction values are drawn from uniform distributions whose expected values are 0, 0.25, 0.5, 0.75, and 1. LV2 uses symmetric lognormal competition: interspecific interaction values are drawn from lognormal distributions with medians 0, 0.25, 0.5, 0.75, and 1, and with `sdlog = 0.5` for the non-zero treatments. In both experiments, diagonal self-interaction is fixed at 1.

The next graph shows the expected distributions of interspecific interaction values specified by these treatments. The zero-competition treatment is a point mass at zero rather than a continuous density, so it is shown as a vertical spike.

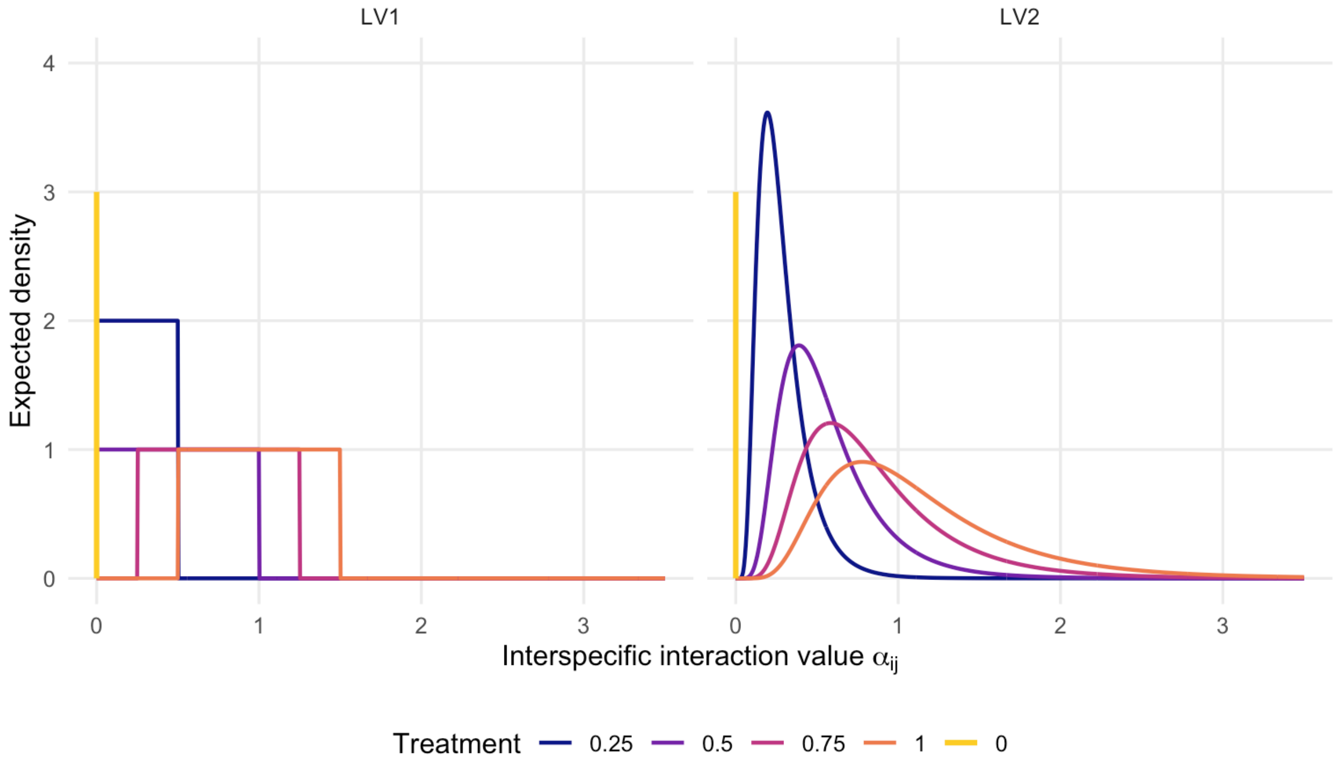


We then calculated the realised value of $p$ for every simulated community in the two LV publication experiments, using the realised interaction matrix and species mean abundances.

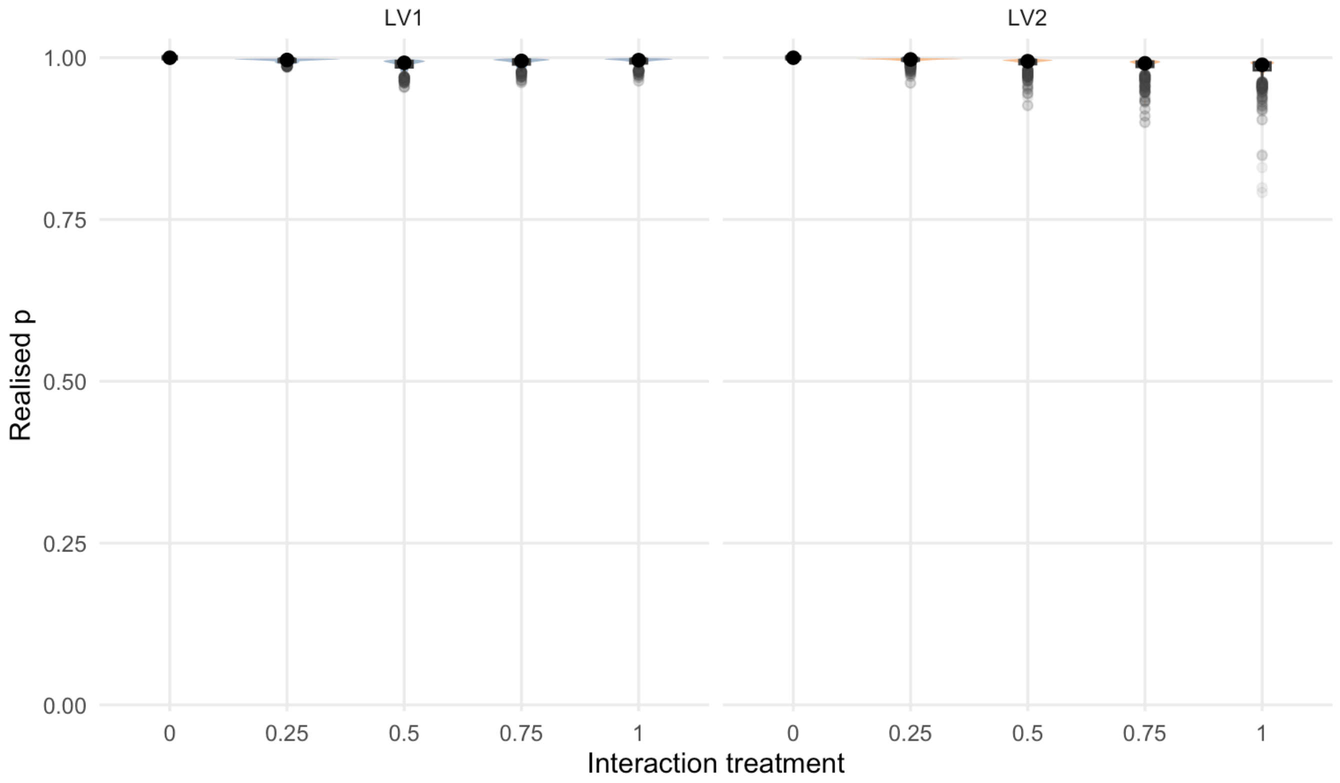


The realised $p$ values are high in both LV experiments. In the no-competition treatment they are $p = 1$. This means that, for most simulated communities, even though interactions can be strong and have large effects on population dynamics, the average interaction structure dominates the pair-specific heterogeneity term in this part of the analytical approximation. The lognormal interaction treatments in LV2 create somewhat lower minimum values of $p$, indicating greater pair-specific heterogeneity in some realised communities, but the median $p$ values remain above 0.9 for all treatments in both experiments. Thus, variation in $p$ alone is not expected to strongly weaken the CP–biomass relationship in these simulations, although the other assumptions listed below still affect how closely the full approximation should hold.

### 10 Scope, assumptions, and limitations

This section states where the derivation should and should not be expected to work. The aim is to make the biological requirements explicit, so that departures in the simulations can be interpreted as informative assumption checks rather than as simple failures of the idea.

1. **Identity-link community performance.** The analysis concerns $CP(t) = \sum_i \tilde{r}_i(T_t)$ and does not derive the binary-viability CP used for the consumer-resource model.
2. **Positive and bounded trajectories.** The log-growth argument requires positive abundances and a negligible finite-time boundary term.
3. **Weak or absent immigration.** Immigration adds the correction in Equation 18.
4. **Timescale separation.** The approximation $\mathbf{N}(t) \approx \mathbf{N}^*[T(t)]$ neglects lags between environmental change and abundance responses.
5. **Feasible and attracting quasi-equilibria.** The temperature-dependent equilibrium must exist, remain non-negative, and be dynamically attracting.
6. **A fixed effective interaction matrix.** The derivation treats $\mathbf{A} = (\beta + \delta)\boldsymbol{\alpha}$ as independent of temperature and time.

7. **Independent pair-specific deviations.** The approximation assumes mean-zero deviations with common variance and neglects correlations or other structured departures from the average interaction matrix.
8. **Ensemble-to-realization approximation.** Terms that vanish across an ensemble need not vanish for a particular finite community.
9. **Second-moment prediction.** The result concerns coefficients of variation and does not describe all features of transient or species-specific dynamics.

### 11 Extension to general link functions

This section places the LV result within the broader community-performance framework. It explains why the derivation applies most directly to the identity-link LV case, and why consumer-resource or viability-based CP measures would need additional assumptions or a separate derivation.

The broader community-performance framework can be written as

$$CP_y(t) = \sum_{i=1}^{m} f_{\text{model},y}\left(P_i[E(t)]\right), \qquad (37)$$

where $P_i(E)$ is a species performance curve and $f_{\text{model},y}$ maps performance onto expected contribution to community variable $y$. The present derivation treats the identity-link LV case,

$$f_{\text{LV},B}(P) = P = \tilde{r}_i(T). \qquad (38)$$

A binary-viability construction corresponds to a nonlinear link such as

$$f(P) = \mathbb{I}(P > 0). \qquad (39)$$

The linear moment mapping used here does not transfer automatically to nonlinear link functions. Extending the theory would require an explicit model connecting transformed performance to abundance, or a suitable local linearization.
